# Germline-encoded V(D)J gene usage does not impose strict constraints on the epitope-specificity of T cell receptors

**DOI:** 10.64898/2026.08.20.745957

**Authors:** Adrian Straub, Yang An, Felix Drost, Kersten Heyer, Zahra Abedi, Monika Hammel, Abagail Delahoussaye, Sabrina Wagner, Anton Mühlbauer, Julian Hönninger, Jack Barton, Karl Moukarzel, Linda Warmuth, Sarah Braun, Laura Valentiner, Corinne Angerpointner, Thomas Pohl, Veit R. Buchholz, Kilian Schober, Matthias T. Warkotsch, Benjamin Schubert, Dirk H. Busch

**Author notes:** These authors contributed equally (Adrian Straub, Yang An).

## Abstract

The theoretical diversity of T cell receptors (TCRs), generated through V(D)J recombination, is enormous, yet the diversity of TCRs capable of recognizing the same epitope remains unknown. Defining this TCR ‘solution space’ is essential for uncovering basic principles that govern TCR specificity. Using single-cell RNA and TCR sequencing, we generated ultra-deep (more than 4000 unique TCRs per epitope) epitope-specific TCR libraries derived from 560 immunized C57BL/6 mice, identifying over 27,000 unique epitope-reactive TCRs across three distinct CD8^+^ T cell epitopes presented by two major histocompatibility complex (MHC) class I alleles. Saturation analyses indicated that the solution space for all studied epitopes comprises many tens of thousands of unique TCRs. Despite highly skewed and peptide-dependent VJ-usage patterns, nearly the entire set of functional germline Vα/β and Jα/β segments was detected at least once within each epitope-specific repertoire. Therefore, diversity of epitope-specific TCRs is not limited by distinct germline combinations, but rather can emerge from a near-to-complete combinatorial space of α- and β-chain, V and J segments paired with compatible CDR3 sequences.

## 1 Introduction

The adaptive T cell immune response relies on the diversity of T cell receptors (TCRs), whose variable regions are generated through somatic recombination of germline-encoded gene segments at the TCRα- and TCRβ-chain loci. While the TCRα locus recombines variable (V) and joining (J) gene segments, the TCRβ locus recombines V and J gene segments alongside one of two available diversity (D) gene segments. Junctional nucleotide deletion and terminal deoxynucleotidyl transferase (TdT)-mediated nucleotide addition at the V-J junction of the α chain and the V-D-J junctions of the β chain generate the hyper-variable complementarity-determining region 3 (CDR3) of each chain ^1,2^. This recombinatorial process produces a vast theoretical sequence space, estimated to exceed 10^15^ unique TCRs, although thymic selection, peripheral homeostasis and space limitations reduce the repertoire diversity of an individual to approximately 10^7^-10^8^ clonotypes ^1,3^. Early studies proposed a division of labor in TCR recognition, in which germline-encoded V segments predominantly contribute to MHC, whereas hyper-variable CDR3 loops predominantly contribute to peptide recognition ^1,5^. Yet, to what extent each component can define or constrain MHC and epitope recognition remains incompletely understood.

Recent advances in single-cell immune repertoire sequencing and high-throughput functional screening have enabled the systematic investigation of TCR repertoires at unprecedented scale. These technologies now allow paired (i.e., α-β) TCR sequences to be linked with antigen specificity and functional phenotypes across large cohorts and experimental systems ^7–12^. Consequently, rapidly growing datasets of antigen-reactive TCRs have transformed our understanding of adaptive immunity. Numerous studies have demonstrated pronounced biases in V, (D), and J segment usage within antigen-specific TCR repertoires, with particular germline segments or combinations strongly enriched in T cell responses to individual epitopes ^13,14^. Hereafter, we refer to these V(D)J combinations, which constitute the fundamental ‘building blocks’ of the TCR, as ‘TCR architecture’ and abbreviate the underlying gene segments *Trav, Trbv, Traj*, and *Trbj* as Vα, Vβ, Jα, and Jβ, respectively. In an extreme example, a malaria-specific response was shown to depend on a particular Vβ, which the authors described as a malaria-specific “immune response gene” ^15^. More recently, Liu et al. introduced TCR specificity profiles building on prevalent germline usage and CDR3 signatures to describe and define epitope-specific TCRs ^16^. Likewise, convergent and “public” TCR sequence features have been well documented ^14,17^. Collectively, these studies suggest a conceptual restriction in TCR architecture in antigen-specific repertoires, while retaining substantial clonotypic diversity, particularly within the CDR3 regions. Nevertheless, no epitope-specific TCR repertoire has yet been sampled close to saturation, and therefore, the full breadth of possible TCR-pMHC interaction modes remains unresolved. Establishing a baseline of TCR datasets that approach saturation of the naturally occurring solution space for defined epitope-specificities is crucial for identifying the sequence and structural determinants of TCR specificity. Deep sequencing of single individuals only yields a limited, highly skewed TCR repertoire dominated by few expanded clonotypes. Thus, sampling many individuals is required to approach the full ‘epitope-specific TCR solution space’. As MHC heterogeneity and other genetic as well as environmental factors can further influence the composition of TCR repertoires ^18–20^, baseline data need first be generated in highly homogeneous cohorts, which is currently best possible with inbred mouse strains kept under standardized housing conditions.

To address the questions: (1) How many distinct TCR solutions can recognize a single defined epitope, (2) how TCR architecture influences T cell function in terms of effective recruitment and clonal expansion, and (3) how TCR architecture constrains epitope-specificity, we deeply sequenced epitope-specific TCR repertories from more than 560 independent C57BL/6 mice, spanning three distinct pMHC specificities. This dataset allowed us to estimate for the first time the overarching space of possible TCR solutions within defined epitope-reactivities and dissect flexibilities in TCR-pMHC recognition patterns in respect to the underlying TCR architecture of epitope-reactive TCRs.

## 2 Results

### 2.1 Generation of saturating epitope-reactive TCR libraries

We first set out to investigate the TCR repertoire in C57BL/6 mice following infection with *Listeria monocytogenes* (L.m.) expressing the Ovalbumin-derived epitope SIINFEKL (OVA; Fig. 1a), combining three complementary approaches to identify OVA-reactive T cells and the corresponding TCR repertoire via single-cell RNA/TCR sequencing. First, TCR downregulation and up-regulation of lysosomal-associated membrane protein 1 (LAMP-1 or CD107a) after antigenic stimulation ^21,22^ (Fig. 1a). Second, H-2K^b^/SIINFEKL pMHC-multimer binding and third, ‘reverse phenotyping’ of CD44^hi^ T cells (this dataset was previously generated for a different study ^23^ and now added to the overall SIINFEKL-specific TCR library (Extended Data Fig. 1a). Beyond the aforementioned criteria, we defined a T cell clonotype as epitope-specific if it was expanded ( ≥2 cells) within a single mouse’s TCR repertoire (‘> threshold’). In total, we analyzed the endogenous OVA-reactive TCR repertoire of 365 C57BL/6 mice, identifying more than 8,600 unique OVA-reactive TCRs (Extended Data Fig. 1b). TCR repertoires captured by the different strategies overlap extensively with one another and with previously collected data, indicating that we robustly captured antigen-reactive TCRs regardless of the isolation approach (Extended Data Fig. 1b).

**Figure 1.**
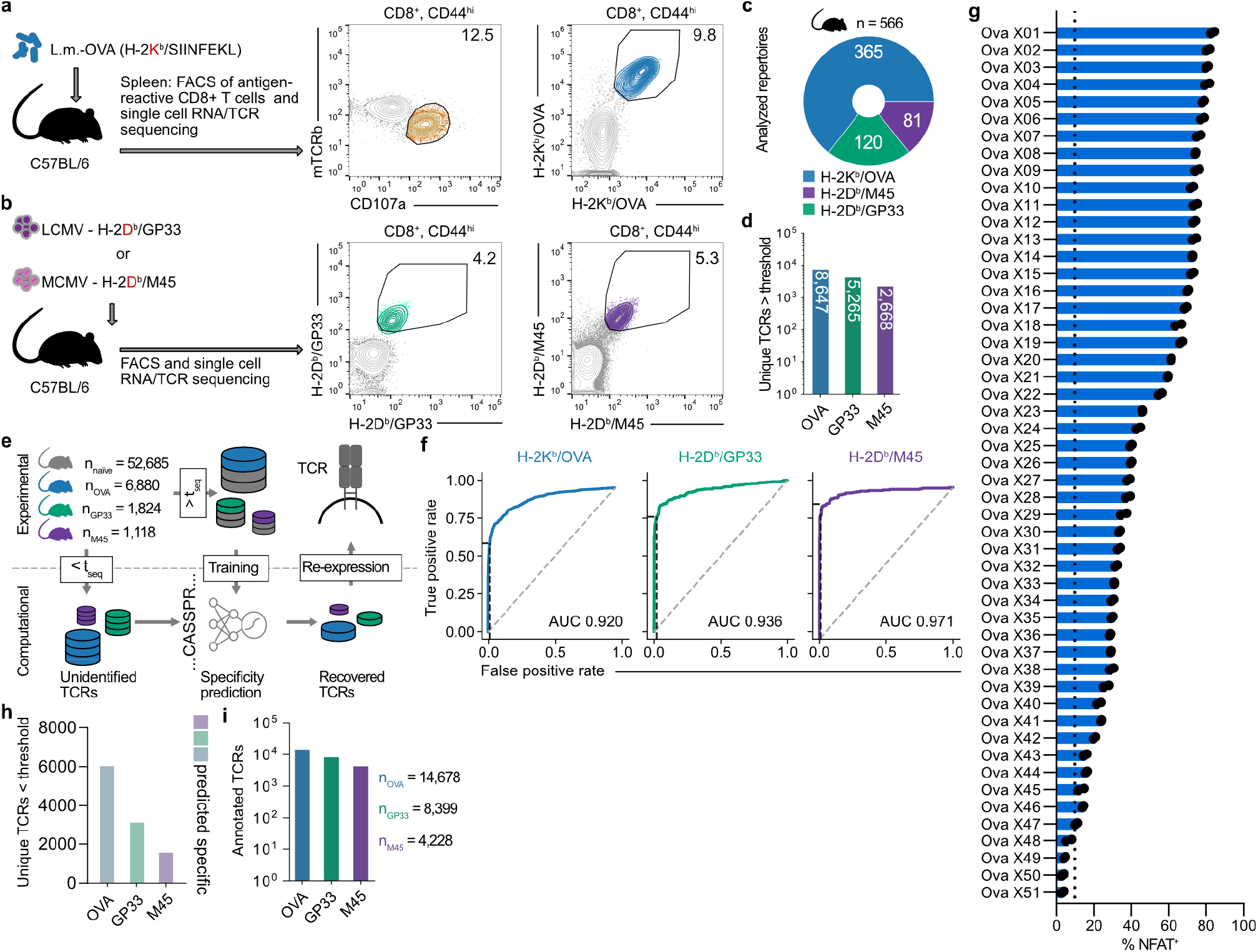
Generation of saturating epitope-specific TCR libraries. (a) C57BL/6 mice were infected with L.m.-SIINFEKL. At day 8 post-infection (p.i.), splenocytes were stimulated with SIINFEKL peptide or left unstimulated. Stimulated antigen-reactive CD8^+^ T cells were identified by CD107a^+^ expression, unstimulated by H-2K^b^/SIINFEKL^+/+^ binding. Representative CD107a^+^ (n = 8 groups of 30 mice) and pMHC^+/+^ (n = 4 groups of 20 mice) stainings are shown. Sorted CD8^+^ T cells were subjected to single-cell-RNA/TCR sequencing. (b) C57BL/6 mice were infected with LCMV or MCMV, at day 8 p.i., splenocytes were subjected to FACS sorting. Antigen-reactive CD8^+^ T cells were identified by pMHC^+/+^ (GP33: H-2D^b^/KAVYNFATC, M45: H-2D^b^/HGIRNASFI) binding. Representative pMHC^+/+^ (n = 6 (GP33), n = 4 (M45) groups of 21 mice) stainings are shown. Sorted T cells were subjected to single-cell-RNA/TCR sequencing. (c) Distribution of mouse TCR repertoires across specificities. (d) Number of unique TCRs across specificities (clone size ≥2). (e) Training of epitope-specific TCR sequence prediction models. (f) Area Under the Receiver Operating Characteristic (AUROC) curve of predictor performances for all specificities on a held-out set, left to right: OVA, GP33, M45. Dashed black line depicts True Positive and False Positive Rate for a threshold of 0.5. (g) NFAT^+^ reporter expression of SIINFEKL-stimulated JTPR engineered with TCRs of the test set. Dotted line indicates NFAT classification threshold. (h) Specific-predicted TCR clonotypes across specificities in the non-expanded (“< threshold”) group. (i) Total number of unique TCR clonotypes across specificities after prediction correction.

To validate the quality of epitope-specific TCR identification, 26 OVA-reactive TCRs were re-expressed in a Jurkat triple parameter reporter (JTPR) cell line. OVA-reactivity upon antigenic stimulation was assessed via activation of the nuclear factor of activated T cells (NFAT) reporter (Extended Data Fig. 1c). Only one of 26 tested TCRs failed to show reactivity to the epitope, corresponding to a false-discovery rate below 5%. Encouraged by these promising results, we generated two additional ultra-deep TCR libraries from 120 and 81 C57BL/6 mice immunized with *lymphocytic choriomeningitis virus* (LCMV) or *murine cytomegalovirus* (MCMV) targeting H-2D^b^/KAVYNFATC (GP33) and H-2D^b^/HGIRNASFI (M45) specific TCRs, respectively (Fig. 1b-c). Overall, we analyzed the antigen-reactive repertoire of 566 C57BL/6 mice, yielding more than 8,600, 5,200, and 2,600 distinct TCR-pMHC pairs targeting H-2K^b^/OVA, H-2D^b^/GP33, and H-2D^b^/M45, respectively (Fig. 1d).

The strict filter of clonal expansion will leave out a larger number of genuinely antigen-reactive TCRs for which only a single T cell had been detected. Therefore, we sought to recover antigen-reactive clonotypes below the expansion threshold and developed an epitope-specific TCR prediction model (Fig. 1e). For each epitope, we trained an ensemble of five models, one per cross-validation split (5.19). Each model used a finetuned ESM-2^24^ to encode the concatenated TCR amino acid sequence, followed by a dense neural network classification head. For negative training data, we profiled the CD8^+^ TCR repertoire of six naïve C57BL/6 mice (Extended Data Fig. 1d). For each cross-validation split, we performed separate hyperparameter tuning, selected the best-performing model on the validation set, and the five resulting models were combined into the final ensemble. This trained ensemble achieved Area Under the Receiver Operating Characteristic (AUROC) scores exceeding 0.90 for all epitopes, with high average precision score (APS) across epitopes (0.80, 0.89, 0.95 for OVA, GP33, M45, respectively) on a held-out test set (Fig. 1f, Extended Data Fig. 1e). To evaluate our model’s predictive accuracy on unseen TCRs, we experimentally expressed 51 TCRs predicted to be epitope-specific from the OVA test set in JTPR cells. We achieved a precision exceeding 90 %, as 47 of 51 candidates showed antigen-dependent NFAT activation after peptide stimulation (Fig. 1g).

In summary, we identified the antigen-reactive TCR repertoire for OVA, GP33, and M45 in over 560 C57BL/6 mice and used this data to establish epitope-specific prediction models that predict antigen-reactivity with high precision. This encouraging result allowed us to confidently recover antigen-reactive TCRs that had previously fallen below our inclusion threshold (Fig. 1h), expanding our ultra-deep TCR library to more than 14,600, 8,300, and 4,200 unique OVA-, GP33-, and M45-reactive TCRs, respectively (Fig. 1i).

### 2.2 The total TCR solution space for functional epitope recognition is remarkably large

We next asked how many distinct TCR solutions can recognize a single defined epitope. Previous studies estimated epitope-specific repertoire sizes within individual hosts but did not address the cumulative diversity across individuals ^25–27^. Chen et al. provided the closest approximation to a population-level epitope-specific repertoire by aggregating deeply sampled CMV- and influenza-specific TCRs across donors, although the total paired TCR solution space remained unresolved ^28^. Across OVA-, GP33- and M45-specific responses, individual mice recruited on average 41, 59, and 55 unique clonotypes, respectively (Fig. 2a). As we sampled additional mice, the cumulative number of distinct clonotypes continued to increase, together with the number of clonotypes shared across animals (Fig. 2b; Extended Data Fig. 2a-b). We calculated the TCR generation probability (P_gen_) using OLGA ^29^ and identified a modest association of publicity with generation probability (*ρ =*0.23, p<0.0001; Fig. 2c; Extended Data Fig. 2a-b).

**Figure 2.**
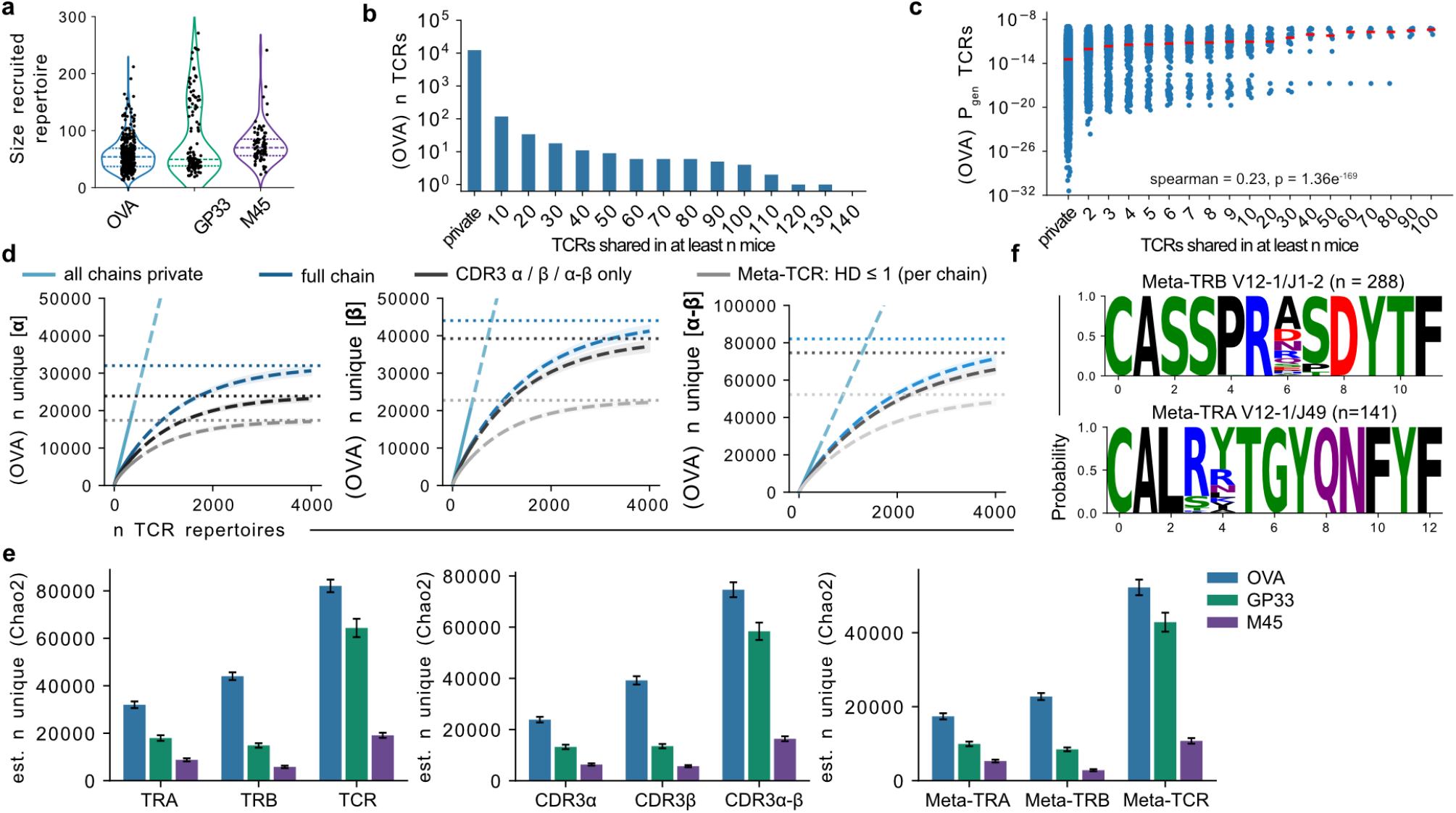
The total TCR solution space for functional epitope recognition is extremely large. (a) Repertoire sizes across specificities; long dashed lines indicate the median and short dashed lines the quartiles. (b) TCR sharedness across donors. (c) Correlation between TCR sharedness and *P*_gen_. (d) Rarefaction and extrapolation curves for unique *α*-, *β*-, and paired *α*-*β*-sequences when sequencing a defined number of repertoires. Blue depicts full chains, dark gray CDR3 only, and light gray Meta-TCRs. The cyan dashed line indicates the expected number of individual sequences if all sequences were private. From left to right: *α, β*, and paired *α*-*β*. HD denotes Hamming distance. Solid lines show the rarefaction curves up to the observed number of repertoires, and dashed lines show the extrapolated curves. Numbers were calculated using iNEXT. Dotted lines show Chao2 asymptotic richness estimates. Shaded areas depict 95% confidence intervals calculated using 100 bootstrap iterations. (e) Estimated total numbers of unique TCRs for different chains and epitopes using Chao2. (f) Logo plots depict CDR3*α* (lower) and CDR3*β* (upper) probability logos of defined Meta-TCRs within a Hamming distance of 1. All error bars depict 95% confidence intervals determined using 100 bootstrap iterations with iNEXT.

The presence of public clones shared across mice allowed us to extrapolate the TCR richness, to quantify the overall TCR “solution space” for a given epitope. TCR richness did not increase linearly with each repertoire (Fig. 2d, cyan, “all private”); instead, the identification of novel unique TCRs followed an asymptotic accumulation curve (Fig. 2d, blue, “full chain”). To derive a lower-bound estimate on total TCR richness, we applied the non-parametric richness estimator Chao2^30^, which infers total diversity from the presence or absence of individual sequences across independently sampled immune repertoires. Both Chao2 and its abundance version Chao1^31^, have previously been used and benchmarked for TCR repertoire diversity profiling ^32–36^. We estimated an asymptotic richness of 82,080 unique OVA-reactive TCRs for paired α-β-chains, comprised of 44,027 unique β-and 31,993 unique α-chains (Fig. 2e). To assess the robustness of these estimates, we applied two complementary diversity estimators, the Incidence-based Coverage Estimator (ICE) ^37^ and the Michaelis-Menten (MM) curve estimator ^38^. Both yielded comparable asymptotic richness estimates (MM: 74,073, Chao2: 82,080, ICE: 96,127; Extended Data Fig. 2e). Notably, estimates of total OVA-reactive TCR richness were similar between defining a clonotype by the full TCR sequence or just by the CDR3 amino acid sequence (Fig. 2e).

Although CDR3 amino acid richness appeared to be extremely large for OVA-reactive TCRs, we identified that many CDR3s differed by only a single amino acid at individual permissive positions without compromising epitope-reactivity. Therefore, we collapsed similar sequences into meta-TCRs: greedy clusters in which a seed CDR3 (sharing identical TCR architecture) absorbs all sequences within a Hamming distance of 1 (Fig. 2f, Extended Data Fig. 2c, Methods 5.21), yielding a smaller, more robustly sampled set with an observed diversity of 11,634 meta-TCRs and an asymptotic estimate of 52,247 using Chao2. Using this meta-TCR definition, we observed an improvement in the sample coverage - the estimated fraction of the total probability mass already captured: meta-TCRs reached a coverage of 0.55, compared to 0.41 for paired TCRs, indicating a more unbiased richness estimate for meta-TCRs (Extended Data Fig. 2f).

We estimated total richness for GP33- and M45-reactive TCRs in the same manner (Extended Data Fig. 2d-e). For GP33 and M45, Chao2 estimated 64,355 and 19,109 unique TCRs, respectively (Fig. 2e). Overall, the estimated TCR solution space was largest for OVA, followed by GP33 and then M45. Notably, the Chao2 estimation of total TCR richness appears linked to the average edit distance of the identified antigen-reactive TCRs (Extended Data Fig. 2g). Across TCRα, TCRβ, or paired TCR chains, M45 and GP33 both ranked below the OVA repertoire in terms of average pairwise edit distance, potentially pointing to a greater permissiveness of the OVA-specific solution space for distinct TCR sequences (Extended Data Fig. 2g).

As comparable studies are missing, this is the first systematic estimate of the population-level TCR solution space within individual epitope specificities. This space is remarkably large, with at least tens of thousands of TCRs for individual epitopes. Within these repertoires, the relative contributions of TCRα- and TCRβ-chain richness to paired-receptor diversity are epitope-dependent.

### 2.3 T cell recruitment is shaped by generation probabilities

Having established that we sampled an unprecedented fraction of the TCR solution space across multiple defined epitope-specificities, we next asked how TCR germline architectures influence recruitment and clonal expansion of T cell clonotypes. First, we analyzed the Vβ repertoire present in antigen-reactive TCRs (Fig. 3a). While the naïve baseline revealed a moderate correlation (*ρ* = 0.47, p = 0.027) between the Vβ frequency and average generation probabilities of TCR β-chains (Extended Data Fig. 3a), Vβ frequencies in epitope-reactive repertoires showed little to no significant correlation, indicating epitope-dependent recruitment of TCR genes among responding clonotypes (Extended Data Fig. 3a).

**Figure 3.**
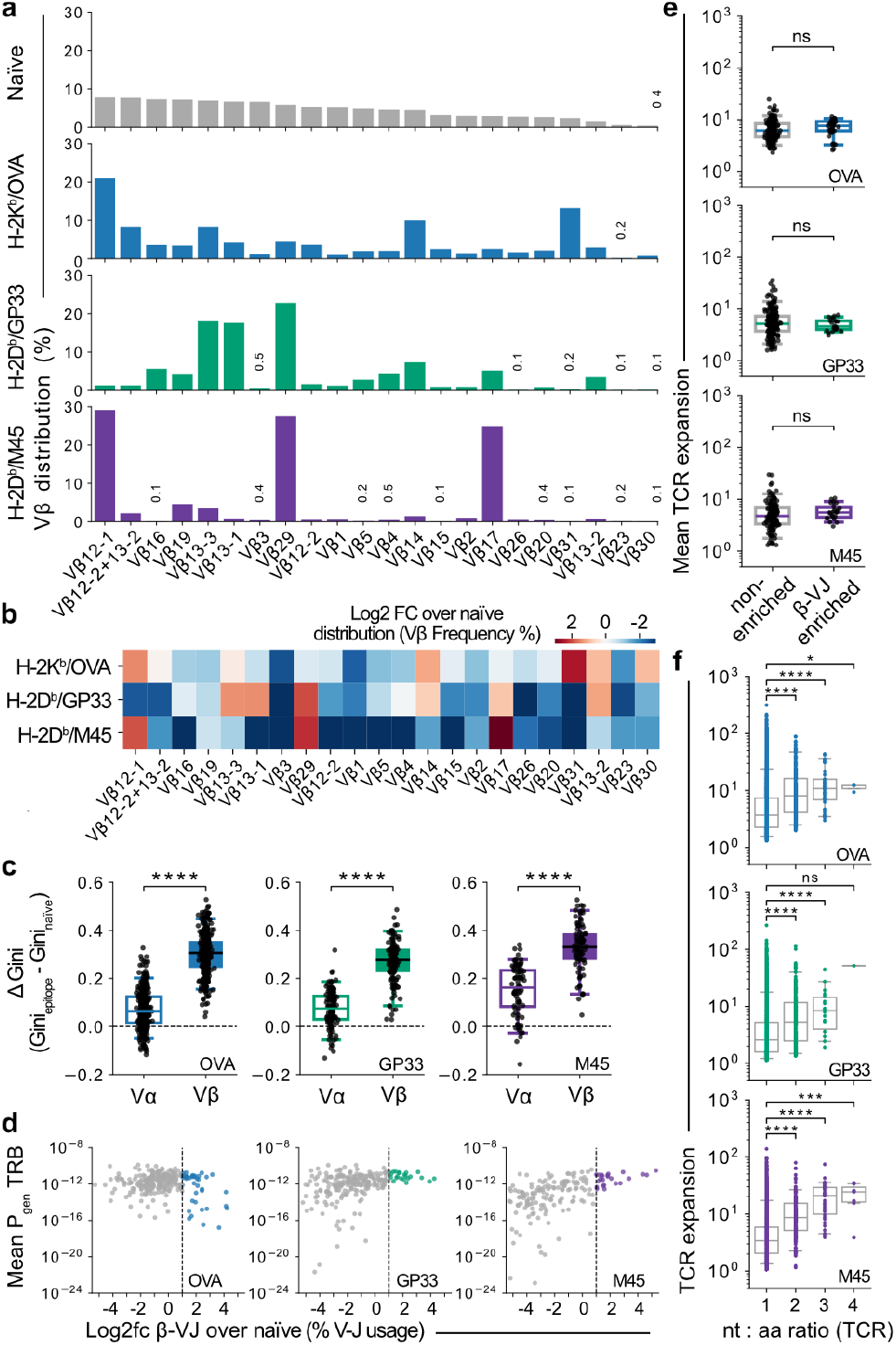
T cell recruitment is shaped by generation probabilities. (a) Frequency of V*β* gene usage in naïve, OVA, GP33 and M45 TCR libraries. (b) Log_2_ V*β* fold change over naïve baseline frequency. (c) Boxplots, delta of Gini coefficient of V*α* and V*β* frequency between epitope-reactive TCRs and naïve baseline. Higher values indicate greater change in skewedness compared to the naïve distribution. Each dot represents the change in Gini coefficient per donor. (d) Identified TCRs were grouped by *β* V-J usage; x-axis depicts *β* V-J log_10_ fold change over naïve baseline, y-axis depicts average *β*-chain Pgen of the respective *β* V-J group. Dotted line indicates a fold change > 2 over naïve as a threshold for enrichment, coloring indicates enriched, gray indicates non-enriched, each dot represents a unique *β* V-J. (e) Boxplots compare average, normalized TCR expansion per *β* V-J group, each dot represents a unique *β* V-J. (f) X-axis depicts nt:aa ratio of aa TCR clonotypes, y-axis depicts expansion, each dot represents a unique TCR. Boxplot whiskers depict 5-95 percentile, in (c) statistical testing was done by a multiple Wilcoxon matched-paired signed rank test with Holm-Šídák correction, all other testing was done using a two-sided multiple Mann-Whitney test with Holm-Šídák correction, * *p* < 0.05, ** *p* < 0.01, *** *p* < 0.001, **** *p* < 0.0001.

We calculated the log_2_ fold change (log2fc) of the Vβ gene frequencies over the naïve baseline and identified between four and five Vβ per epitope to be specifically enriched (log2fc > 1; Fig. 3b). Complementarily, we assessed Vα gene frequencies and calculated the Gini coefficient for Vβ and Vα genes in each epitope-reactive repertoire, as well as the naïve baseline. We identified a significantly stronger change in skewness (ΔGini) for Vβ, compared to Vα (Fig. 3c), suggesting that the Vβ germline contributes more strongly to epitope-specific selection in T cell recruitment than Vα. Consequently, we then focused on the TCRβ germline architecture in more detail. We calculated the log2fc of every detected β V-J combination over the naïve baseline and defined a specific architecture to be enriched if its log2fc exceeded 1 (Fig. 3d) and asked whether preferred architectures translate to a larger clonotypic response *in vivo*. Remarkably, enriched and non-enriched TCR architectures showed comparable clonal expansion *in vivo* (Fig. 3e), suggesting that preferential TCR gene usage does not determine effective recruitment or the magnitude of the clonal response.

Additionally, the β V-J enrichment in the OVA-reactive repertoire was not associated with β-chain generation probability. In contrast, enriched β-chain architectures in the GP33- and M45-reactive repertoires contained a significantly higher proportion of high-*P*_*gen*_ chains (Extended Data Fig. 3b). Previous work suggested that biases in TCR generation may favor receptors capable of recognizing foreign antigens ^39^. We therefore stratified clonotypes by the median into high- and low-*P*_*gen*_ groups (Extended Data Fig. 3c). At the individual clonotype level, however, generation probability did not predict expansion capacity, as high- and low-*P*_*gen*_ TCRs reached comparable response sizes (Extended Data Fig. 3c).

Alternatively, we have previously hypothesized that the association between *P*_*gen*_ and clonal abundance could be explained by repeated generation of identical TCRs during thymocyte maturation, resulting in larger naïve precursor pools ^38^. While we previously inferred this potential mechanism from large-scale unpaired repertoire analyses, the generated paired single-cell dataset in this study allowed us to directly relate TCR *P*_*gen*_ to the expansion of individual epitope-reactive clonotypes. Overall, TCR *P*_*gen*_ positively correlated with the nucleotide-to-amino acid (nt:aa) ratio of clonotypes shared across mice (Extended Data Fig. 3d). We used the nt:aa ratio within individual mice as a proxy for convergent recombination and inferred naïve precursor frequency. Clonotypes encoded by multiple nucleotide rearrangements (nt:aa > 1) underwent significantly greater expansion than those represented by a single nucleotide sequence (Fig. 3f).

Together, these findings support a model in which TCR generation probability shapes T cell recruitment by increasing the likelihood of repeated receptor generation and thereby favoring its representation in a precursor pool, rather than by conferring an epitope-specific proliferative advantage to individual T cell clones. In contrast, a specific β V-J architecture did not contribute to stronger proliferation.

### 2.4 TCR germline architecture enables higher permissiveness in epitope-specific TCR features

Numerous studies have documented epitope-dependent biases in TCRβ gene usage following infection, vaccination or in tumor responses using either pMHC tetramers ^40^ or anti-Vβ antibodies ^41^. Thus, we asked how TCR germline architectures shape epitope-specific CDR3β sequence motifs by comparing enriched and non-enriched architectures. To focus the analysis on the hypervariable junctional region of the CDR3, we trimmed two C-terminal and three N-terminal amino acids and enumerated all contiguous 2- and 3-mer motifs within the remaining CDR3β sequence. K-mer diversity was quantified for each unique TCRβ V-J combination using the Miller-Madow sample-size-corrected Shannon entropy of the observed k-mer distribution (H_MM_; Methods 5.18). Both k-mer distributions displayed significantly higher entropy among preferred architectures (Fig. 4a; Extended Data Fig. 4a), pointing to higher restriction in motifs of non-enriched architectures. Additionally, across all epitope specificities, enriched TCRβ V-J combinations displayed significantly greater amino-acid diversity per CDR3β position than non-enriched combinations, indicating a broader range of junctional sequence solutions (Fig. 4b; Methods 5.18). Although all TCRβ architectures exhibited conserved patterns of CDR3β amino-acids relative to the naïve baseline, these patterns were more pronounced in non-enriched architectures, which showed stronger position-specific amino-acid preferences and reduced junctional sequence flexibility compared with enriched architectures (Fig. 4a-c; Extended Data Fig. 4a-b).

**Figure 4.**
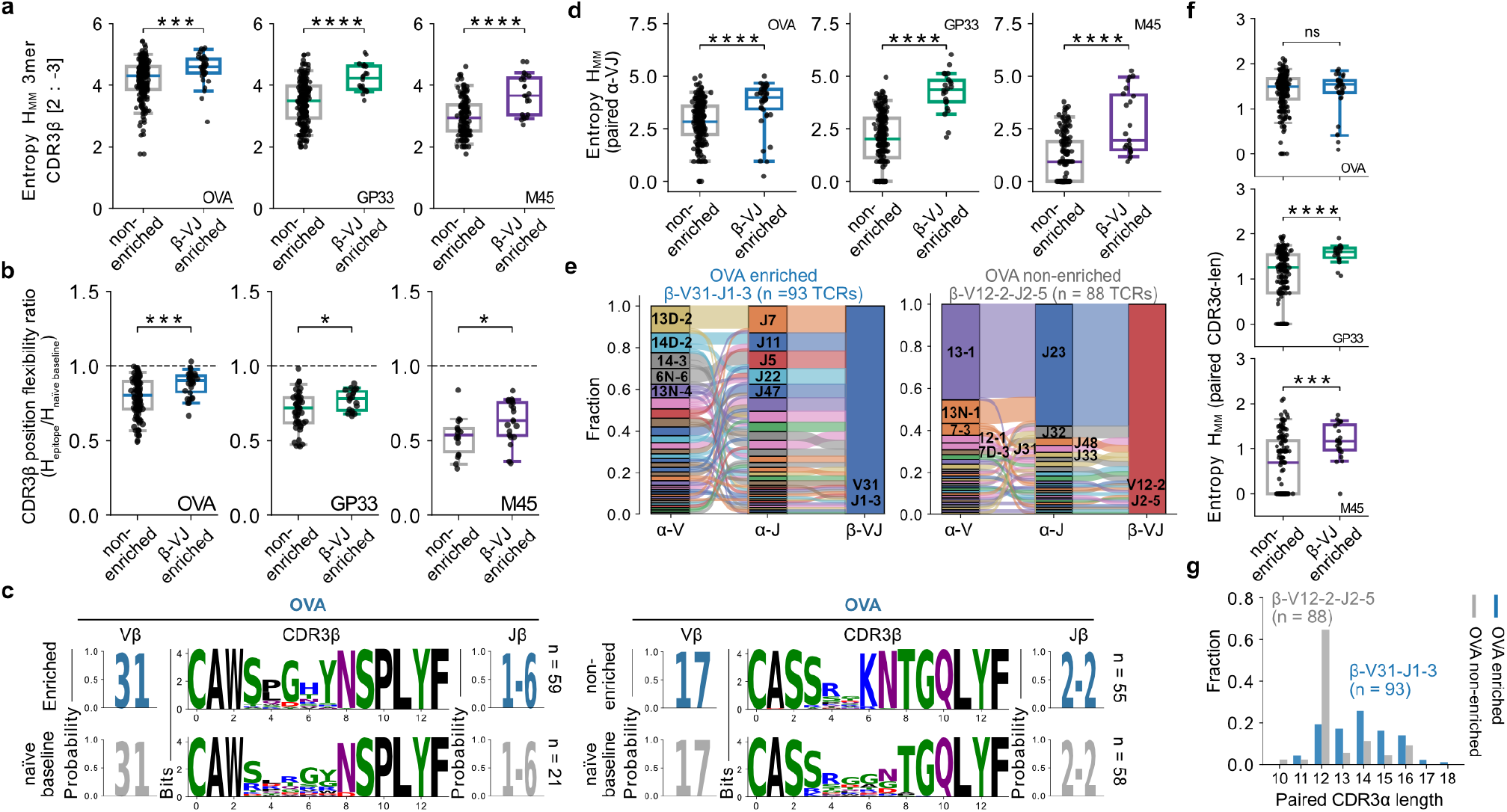
TCR germline architecture enables higher permissiveness in epitope-specific TCR features. (a) H_MM_ of CDR3 kmers (k=3) of enriched and non-enriched *β* V-J; each dot represents a unique *β* V-J. (b) Ratio of H_MM_ per CDR3*β* position between enriched and non-enriched *β* V-J over *β* V-J-matched naïve baseline. Ratio > 1 means higher entropy compared to baseline, ratio < 1 means lower entropy compared to baseline. (c) CDR3*β* logo plots of OVA-reactive TCRs and naïve baseline, left: probability logo of V*β*, right: probability logo of J*β*. (d) H_MM_ of paired *α* V-J usage between non-enriched and enriched *β* V-J groups, each dot represents H_MM_ of a unique *β* V-J. (e) *α* V-J-*β* V-J ribbon plots of OVA-reactive TCRs. The displayed *β* V-J group was selected for comparable size within IQR (0.35 - 0.65) of calculated H_MM_ displayed in (d). (f) H_MM_ of paired CDR3*α*-length of non-enriched and enriched *β* V-J groups; each dot represents H_MM_ of a unique *β* V-J. (g) CDR3*α* length spectratypes of OVA-reactive TCRs from (e) (gray, non-enriched, blue, enriched). Boxplot whiskers depict 5-95 percentile; all statistical testing was done using a two-sided multiple Mann-Whitney test with Holm-Šídák correction. * *p* < 0.05, ** *p* < 0.01, *** *p* < 0.001, **** *p* < 0.0001.

Building on these observations, we asked whether preferred TCR germline architectures permit greater flexibility in α-β-chain pairing while preserving epitope recognition. We quantified the diversity of the paired α V-J usage for each unique β V-J architecture using H_MM_. Enriched TCRβ V-J architectures displayed significantly greater diversity of corresponding α V-J pairings than non-enriched architectures, indicating that preferred β-chain architectures can accommodate a broader range of complementary α-chain solutions (Fig. 4d). Specifically, enriched TCRβ V-J combinations showed considerable flexibility in both paired Vα and Jα usage (Fig. 4e). In contrast, the non-enriched fraction showed stronger restriction on the paired Jα, as well as Vα segments (Fig. 4e, Extended Data Fig. 4c).

Along these lines, the preferred TCRβ architecture for GP33- and M45-reactive TCRs displayed significantly larger flexibility in the paired CDR3α length (Fig. 4f-g; Extended Data Fig. 4c). Although, this effect was not significant for OVA-reactive TCRs, 35 out of 39 enriched TCRβ V-J combinations tolerated diverse CDR3α length pairings (Fig. 4f-g). Analogously, we investigated the enrichment and entropy profile for α V-J architectures. The observed effect in TCR chain pairing permissiveness was more pronounced for preferred TCRβ V-J genes, which displayed more flexibility in V-J and CDR3 length pairing partners compared to enriched α V-J groups (Extended Data Fig. 4d).

Overall, our findings support the hypothesis that preferred TCRβ germline architectures contribute to epitope recognition and therefore allow for stronger permissiveness in complementary chain pairing partners and usage of flexible CDR3β motifs.

### 2.5 Almost all TCR germline architectures can give rise to a highly functional epitope-reactive TCR

Although specific TCR germline architectures were consistently favored during clonotype recruitment across all epitope-specific repertoires, they were not associated with increased clonal expansion. This prompted us to ask whether TCR architecture primarily constrains epitope specificity, if not functionality. Remarkably, we identified across H-2K^b^/OVA-, H-2D^b^/GP33-, and H-2D^b^/M45-reactive TCRs all naturally occurring Vβ, Jβ, as well as Jα gene segments, alongside close to all Vα gene segments (Fig. 5a). Furthermore, we found that essentially every Vβ segment was capable of contributing to an epitope-specific response, indicating broad accessibility of the TCRβ germline repertoire during epitope-specific recruitment. For OVA, GP33, and M45, respectively, nearly any Vβ and 96%, 82%, and 75% of observed β V-J combinations contained clonotypes detected at least five times after infection (Fig. 5b; Extended Data Fig. 5a). This broad recruitment was also evident at the level of germline architecture coverage: we identified more than 95%, 85%, and 62% of all possible β V-J combinations within the OVA-, GP33-, and M45-specific repertoires, respectively (Fig. 5c). Thus, despite strong differences in the relative abundance of individual architectures, a single epitope-specific repertoire displayed near complete recruitment of the available TCRβ germline space.

**Figure 5.**
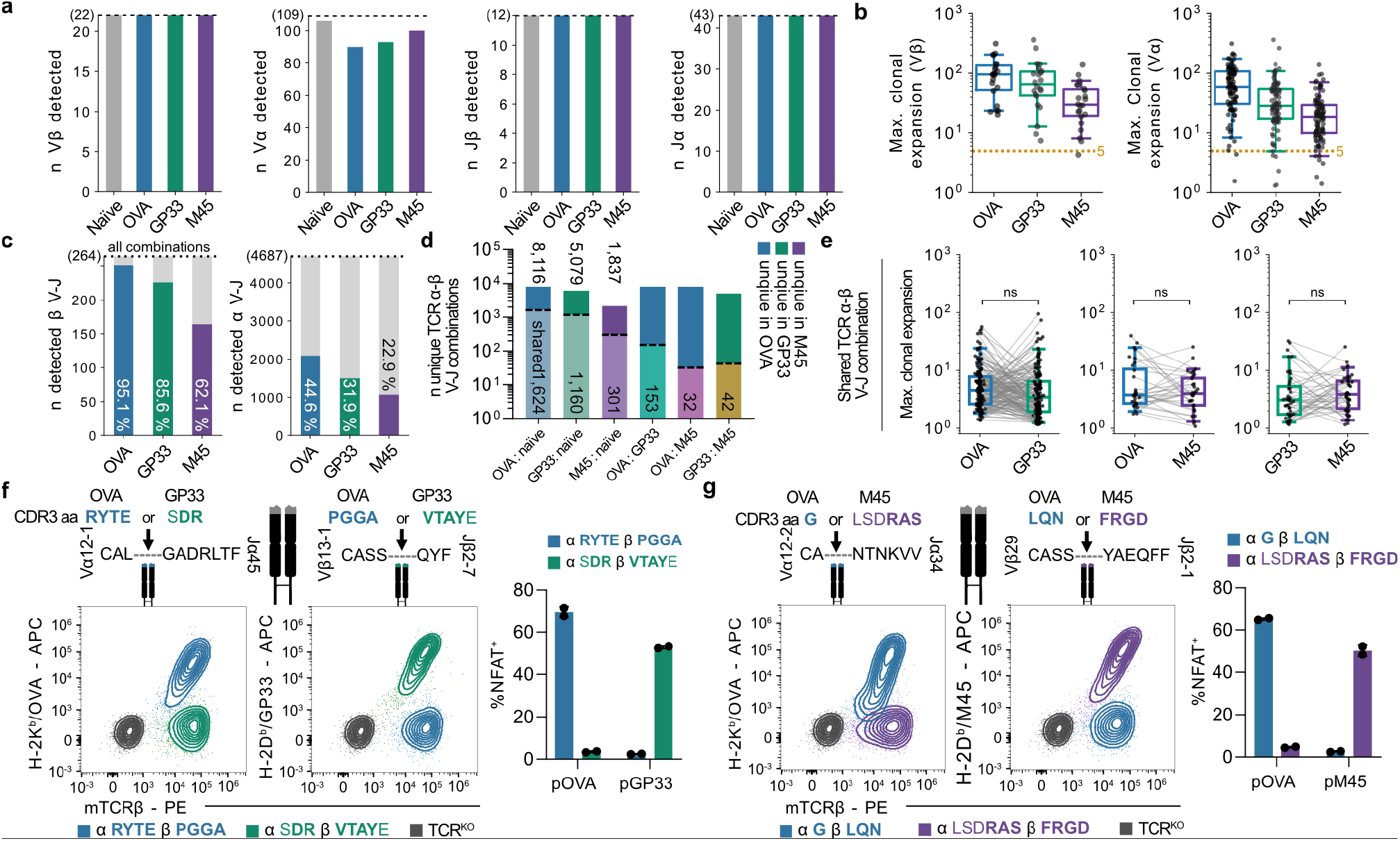
Almost all TCR germline architectures can give rise to a highly functional epitope-reactive TCR. (a) Identified V*β*, V*α*, J*β* and J*α* genes in each TCR repertoire. (b) Maximum detected clonal expansion of a TCR per V*β* (left) and V*α* gene (right). Dotted line indicates expansion ≥5 (c) All detected *β* V-J (left) and *α* V-J (right) combinations. Dotted line indicates total possible combinations. Numbers (white) indicate the percentage detected of total combinations. (d) Unique or shared TCR *α*-*β* V-J combinations between epitope-reactive TCRs or naïve baseline. Upper bars indicate unique TCR *α*-*β* V-J combinations, lower bars indicate shared combinations with the indicated group. Numbers above indicate detected unique combinations per epitope. (e) Maximum clonal expansion of TCRs with TCR *α*-*β* V-J combination shared between depicted groups, each dot represents the max. observed expansion of a unique TCR *α*-*β* V-J combination, connected lines indicate identical TCR *α*-*β* V-J. (f) Identical TCR *α*-*β* germline architecture recognize two distinct epitopes (OVA and GP33), colored amino acid letters indicate minimal exchange of non-templated CDR3 in the *α* and *β* locus to switch functional epitope recognition to the respective epitope. Contour plots display TCR expression (x-axis) and multimer binding of TCR-engineered JTPR; coloring indicates epitope-reactivity. Barplot depicts antigenic stimulation of JTPR; coloring indicates the same TCR, x-axis indicates antigenic peptide. (g), as in (f) but for TCRs with identical germline directed against OVA and M45. Boxplot whiskers depict 5-95 percentile; all statistical testing was done using a two-sided multiple Mann-Whitney test with Holm-Šídák correction, except (F); where statistical testing was done by a multiple Wilcoxon matched-paired signed rank test with Holm-Šídák correction. * *p* < 0.05, ** *p* < 0.01, *** *p* < 0.001, **** *p* < 0.0001.

A similarly broad pattern was observed for the TCRα chain. More than 97%, 94%, and 93% of Vα segments detected in the OVA-, GP33-, and M45-reactive repertoires, respectively, contributed to clonotypes observed at least five times (Fig. 5b). At the level of complete α V-J architectures, 68%, 59%, and 50% contained substantially explaned clonotypes (expansion ≥ 5; Extended Data Fig. 5a), while overall α V-J recruitment encompassed 44%, 31%, and 22% of the possible combinations for OVA, GP33, and M45, respectively (Fig. 5c). When considering both chains together, the OVA-specific repertoire alone recruited more than 0.7% of all possible combined TCRαβ V-J germline architectures (Extended Data Fig. 5b).

Following these observations, we investigated whether the overall TCRαβ germline architecture would be unique to a given epitope-specificity. We identified extensive overlap in TCR germline architectures between any epitope-specificity and the naïve baseline (> 1,600; > 1,100; > 300 TCRαβ V-J combinations in OVA, GP33, and M45, respectively), which should theoretically incorporate any epitope-specificity. Additionally, more than 150, 30, and 40 TCRαβ V-J combinations were shared between OVA and GP33, OVA and M45, and GP33 and M45, respectively (Fig. 5d).

Remarkably, when we compared the response size of these TCRs, any of these shared germline architectures constituted a clonotype which was in most cases identified to be clonally expanded, independent of its epitope-specificity (Fig. 5e). Furthermore, we did not identify significant differences in the maximal TCR response size when the same TCR germline architecture was utilized in either the one or the other antigen-reactive T cell response (Fig. 5e).

To validate these findings, we selected two unique TCRαβ germline architectures shared either between H-2K^*b*^/OVA and H-2D^*b*^/GP33 (Vα12-1 Jα45 + Vβ13-1 Jβ2-7), or H-2K^*b*^/OVA and H-2D^*b*^/M45 (Vα12-2 Jα34 + Vβ29 Jβ2-1) for re-expression in JTPR cells. Notably, for the selected Vα12-1 Jα45 + Vβ13-1 Jβ2-7 TCR germline, i.e., fully identical TCR sequence besides a minor amino acid exchange in the CDR3 of the α-chain (OVA: RYTE instead of GP33: SDR) and β chain (OVA: PGGA instead of GP33: VTAYE) led to loss of binding of H-2D^b^GP33 and maintenance of binding only to H-2K^b^/OVA and vice versa (Fig. 5f). Complementary, we performed antigenic stimulation and observed only functional recognition directed against a single epitope (Fig. 5f). The same outcome was observed, when we investigated the TCR Vα12-2 Jα34 + Vβ29 Jβ2-1 germline usage in respect to reactivity to H-2K^b^/OVA and H-2K^b^/M45. Replacing “OVA: G” with “M45: LSDRAS” for the α-chain, as well as “OVA: LQN” with “M45: FRGD”, led to single binding of H-2K^b^/OVA (and vice versa of H-2K^b^/M45), as well as functional recognition of only a single epitope after antigenic stimulation (Fig. 5g).

In conclusion, a broad range of germline architectures can support recognition of the same epitope. We next asked whether epitope-specific CDR3 motifs are similarly reusable across distinct germline architectures while preserving specificity. We identified significantly enriched 3-, 4-, and 5-mer motifs within the CDR3β regions of epitope-specific TCRs relative to the naïve baseline (Extended Data Fig. 5c). While many epitope-associated k-mers were preferentially associated with a dominant Vβ segment, identical motifs could be incorporated into multiple distinct β V-J architectures (Extended Data Fig. 5d-e). Importantly, TCRs incorporating the same motif through less frequently used Vβ segments could still give rise to expanded epitope-specific T cell clones (Extended Data Fig. 5f).

Overall, these data demonstrate - at least for the three epitopes examined here at unprecedented resolution - that a specific TCR germline architecture can favor the generation of a highly functional TCR within epitope-specific repertoires. Nonetheless, the underlying TCR architecture is not imposed as an absolute requirement for epitope recognition and can be shared even between repertoires restricted by different MHC alleles. At the same time, highly similar epitope-specific CDR3 configurations are equally reusable across different architectures.

## 3 Discussion

A central open question in T cell immunology is to what extent functional TCR-pMHC recognition is, next to CDR3 regions, dependent on distinct TCR germline architectures. Early work established a canonical model in which germline-encoded CDR1/2 loops predominantly contact MHC, whereas CDR3 loops predominantly engage peptide ^1,5,6^. Subsequent structural and functional studies identified recurrent germline-encoded TCR-MHC contacts and V gene-dependent docking preferences ^42–44^, while repertoire-scale analyses revealed strong epitope-associated biases in V-J usage ^16^. However, this division is not absolute: germline-encoded loops can contribute in some cases directly to peptide recognition, whereas CDR3 regions can contribute to MHC binding as well ^45–47^. TCR specificity is therefore increasingly viewed as a cooperative property of the complete TCR-pMHC interface ^48^. It remains unresolved whether preferred TCR germline architectures exist that are required for functional epitope recognition or rather increase the likelihood that an epitope-compatible CDR3 solution can emerge. By deeply sampling H-2K^b^/OVA-, H-2D^b^/GP33-, and H-2D^b^/M45-specific TCR repertoires across more than 560 C57BL/6 mice, we show in this report that defined epitope specificities can be recognized by at least tens of thousands of TCR solutions basically spanning the entire architectural TCR germline space. Preferred germlines exist, but they increase permissiveness for CDR3 sequence and chain pairing diversity rather than acting as strict determinants of specificity.

Previous estimates of epitope-specific TCR diversity were largely derived from individual hosts and limited sampling depths and therefore captured only a fraction of the receptors capable of recognizing a given pMHC. Paired TCR chain analysis progressively revealed substantially greater diversity, including large individualized repertoires directed against single viral epitopes ^14,28,49^. Our findings extend this concept for the first time from individual repertoire diversity to the large-scale inter-individual population level, revealing that functional recognition of a single pMHC is compatible with an unexpectedly large number of distinct TCR germline architectures and CDR3 solutions. Importantly, this diversity is not randomly distributed. Extensive repertoire breadth coexists with reproducible biases in germline architecture and CDR3 composition ^13,14,28^. Thus, functional epitope recognition appears highly permissive, accommodating many interaction possibilities within an epitope-specific solution space.

Notably, the enrichment of particular germline segments did not translate into greater proliferative capacity after recruitment, as enriched and non-enriched TCRβ V-J architectures displayed comparable clonal expansion. Instead, the response size associates with TCR generation probability. Receptors with high TCR P_gen_ can arise repeatedly through independent thymic rearrangement events and are therefore more likely to occur at increased frequency in the naïve repertoire and to be shared between individuals ^38,50–52^. Together, our findings suggest that TCR germline architecture influences the likelihood with which functional epitope recognition can be achieved ^53,54^, while generation probabilities predominantly shape the numerical representation of clonotypes before antigen encounter. Consequently, clonotype abundance after immunization can also reflect differences in precursor frequency, and is not necessarily driven by intrinsic properties of the TCR, such as superior receptor avidity.

TCR architectures enriched within epitope-specific repertoires exhibited greater CDR3 diversity and increased *α* − *β*-chain pairing flexibility than non-enriched architectures. Previous studies reported biased antiviral repertoires, constrained CDR3 motifs, and strong V-J enrichment, which can lead to the interpretation that individual pMHCs preferentially recruit restricted receptor solutions ^13,14,16,28^. However, repertoire bias does not necessarily imply limited sequence-level diversity: preferred V-gene usage can coexist with substantial CDR3 variation ^55^. Structural studies provide a potential mechanistic basis for this observation. Germline-encoded TCR-MHC interactions can favor particular docking geometries, while CDR3 sequences contribute to recognition not only through direct peptide contacts but also by shaping how germline-encoded elements interact with the pMHC complex within the same architecture ^42,43,53,54^. Following this rationale, favorable germline architectures would be expected to accommodate a broader range of CDR3 solutions rather than constrain them. Our data provide experimental support for this hypothesis, suggesting that enriched germline architectures act less as restrictive templates and more as permissive recognition modes capable of supporting a larger spectrum of compatible CDR3 configurations.

Importantly, we detect near-complete recruitment of all TCR V and J architectures across all three epitope-reactive repertoires. The broad sharing of TCR germline architectures across epitope-specific repertoires, indicates that architectural enrichment does neither reflect an absolute requirement for recognition, nor that there is a strict peptide- or MHC-specific restriction at the level of individual TCR germline gene segments. The observation, that the overall combinatorial space of β and α V-J architectures exceeds a remarkable 95%, and 44%, respectively, of the total possible space within OVA-specific TCRs alone, suggests that this finding is not by coincidence unique to the here addressed antigen-reactivities. Particular germline combinations have been proposed to provide reusable recognition architectures whose eventual specificity is resolved through the CDR3 sequence and *α* − *β*-chain pairing ^46,56,57^. Along the lines of these studies, we observe and add to, that identical TCR germline architectures, which translate to a near-identical amino acid sequence of a TCR, can participate with minor changes in CDR3 regions in multiple epitope-specific repertoires, including epitopes presented by distinct MHC alleles.

Thus, TCR-pMHC recognition may be better understood as a probabilistic ‘compatibility landscape’ in which germline architectures shape the likelihood of productive solutions, while the CDR3 sequence ultimately specifies and redirects pMHC recognition.

Structural studies have suggested that germline-encoded CDR1/2 interactions with the MHC helices help position the TCR over the pMHC and constrain functional docking geometries ^42,43,58^. Our data suggest such constraints don’t apply as a ‘general rule’. As both the CDR3 and germline architecture are remarkably flexible within the same epitope context, one could argue that the natural V(D)J-diversity may be an ‘optimized’ driver of CDR3 diversity, instead of leaving the hyper-variable CDR3 generation to pure chance of nucleotide additions and deletions. Current pan-epitope TCR-specificity prediction models remain unable to reliably generalize to unseen epitopes ^59,60^, potentially because the available training datasets capture only the tip of the iceberg of the remarkably large TCR-pMHC recognition space.

In summary, we observe that firstly, a huge variety of TCR germline architectures can participate in recognizing the same epitope, secondly, identical architectures can be used to identify different epitopes, even across MHC context and thirdly, the same CDR3 motif can be utilized across different TCR germline architectures, while preserving the same epitope specificity. Together, our findings suggest a general organizing principle of TCR recognition, in which TCR germline architectures provide a reusable scaffold with unique likelihoods of generating a CDR3 configuration which defines the specific pMHC recognition. Our novel data provide a fundamental baseline for understanding and, ultimately, engineering TCR specificity by leveraging the reusable nature of germline architectures.

## ACKNOWLEDGEMENTS

We thank members of the Busch lab for their support in this research. We thank A. Hochholzer for technical help and generation of pMHC-multimers. We thank I. Andrä for support in cell sorting. The ultra-deep TCR libraries were created as part of the MATCHMAKERS team, supported by the Cancer Grand Challenges partnership financed by CRUK (CGCATF-2023/100002) and the National Cancer Institute (1OT2CA297204-01). A.S., K.H., Z.A., M.T.W. and D.H.B are members of the MATCHMAKERS team. The work was further funded by SFB-TRR 338/1 2021-452881907 project A01 (D.H.B.), Deutsche Krebshilfe 70113918 (V.R.B.), European Research Council starting Grant 949719, SCIMAP, (V.R.B.) and the BMBF grant DeepTCR #031L0290A (B.S.). A.Y. and F.D. are supported by the Helmholtz Association under the joint research school “Munich School for Data Science - MUDS”. F.D. acknowledges financial support from the Joachim Herz Stiftung.

## AUTHOR CONTRIBUTIONS

Conceptualization: A.S., D.H.B.

Investigation: A.S., Y.A., F.D., K.H, A.D., Z.A., M.T.W., K.S., M.H., S.W., A.M., J.H., J.B., K.M., L.W., S.B., L.V., C.A., T.P.

Visualization: A.S., Y.A., F.D., K.H, A.D., Z.A., M.T.W.

Supervision: A.S., B.S., M.T.W., D.H.B.

Writing – Original Draft: A.S., Y.A., F.D., K.H., M.T.W., K.S., B.S., D.H.B.

Writing – Review and Editing: A.S., Y.A., F.D., K.H., M.T.W., K.S., B.S., D.H.B.

Funding Acquisition: B.S., D.H.B

## Resource availability

Further information and requests for resources and reagents should be directed to and will be managed by the lead contact.

## Data Availability

Data are made available from the lead contact upon reasonable request.

## Code Availability

Code for generating the epitope-specific prediction models is made available upon reasonable request. Any additional information and notebooks containing all steps of data processing and analysis of single-cell RNA sequencing reported in this paper are available from the lead contact upon reasonable request.

## Materials availability

This study did not generate new unique reagents.

## 4 Extended data

**Extended data 1.**
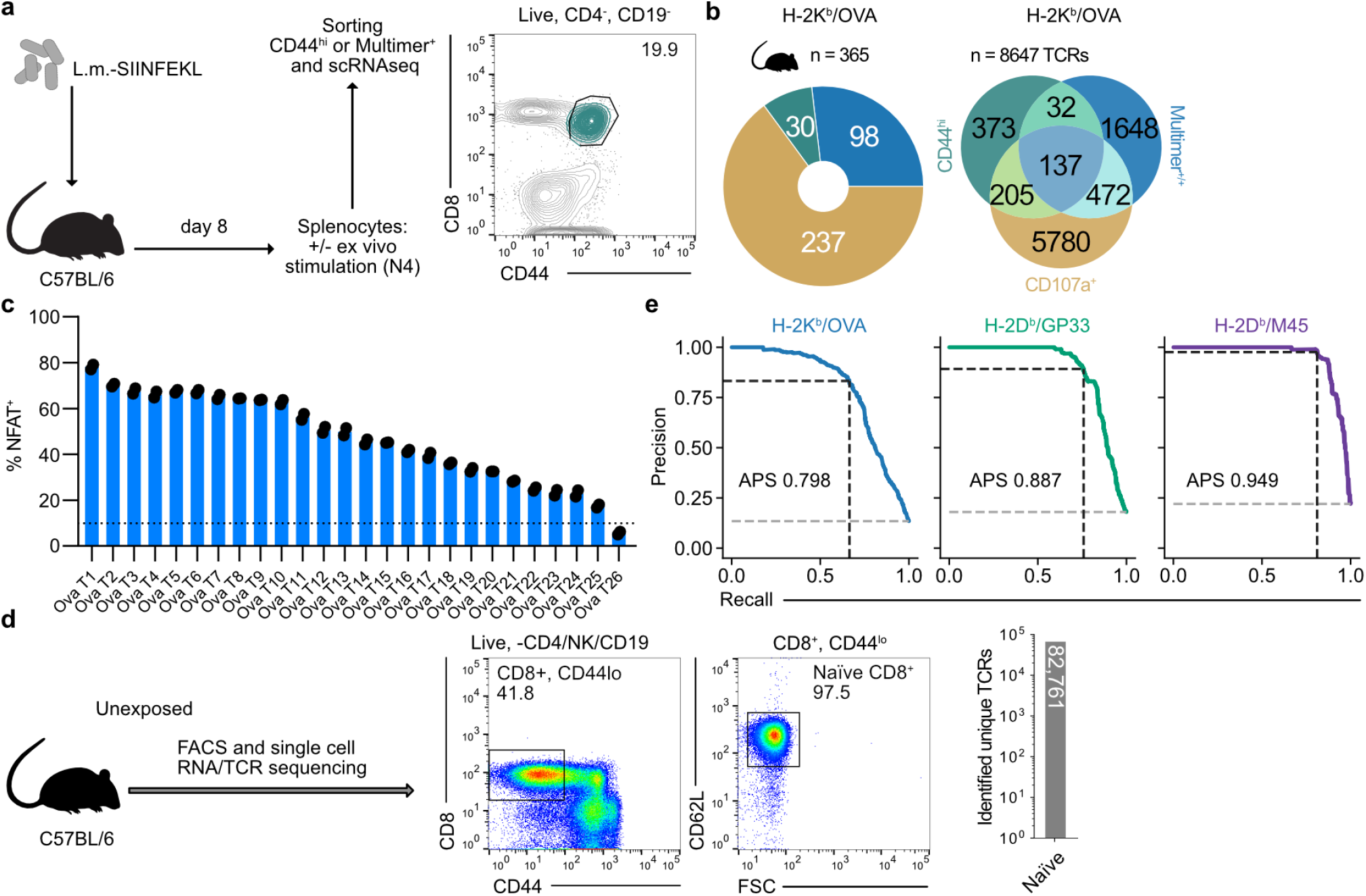
related to Figure 1: Generation of saturating epitope-specific TCR libraries. (a) Schematic: Data acquisition (*n* = 30 mice) of previous H-2K_*b*_/OVA reactive TCR library via reverse phenotyping (Straub et al., 2023) appended to our data analysis. (b). Left: Data distribution of analyzed OVA-reactive TCR repertoires by different isolation methods and right: overlap of OVA TCR sequences across experiments and isolation methods. (c) NFAT^+^ reporter signal expression of SIINFEKL (10^−5^ M) stimulated Jurkat triple parameter reporter (JTPR) cells engineered with SIINFEKL library TCRs of the training set of the predictor. (d) Naïve CD8 T cells from six unexposed C57BL/6 mice were subjected to single-cell-RNA/TCR sequencing. Representative FACS plots (*n* = 6) are shown. (e) APSs of predictor performances for all specificities, left to right: OVA, GP33, M45.

**Extended data 2.**
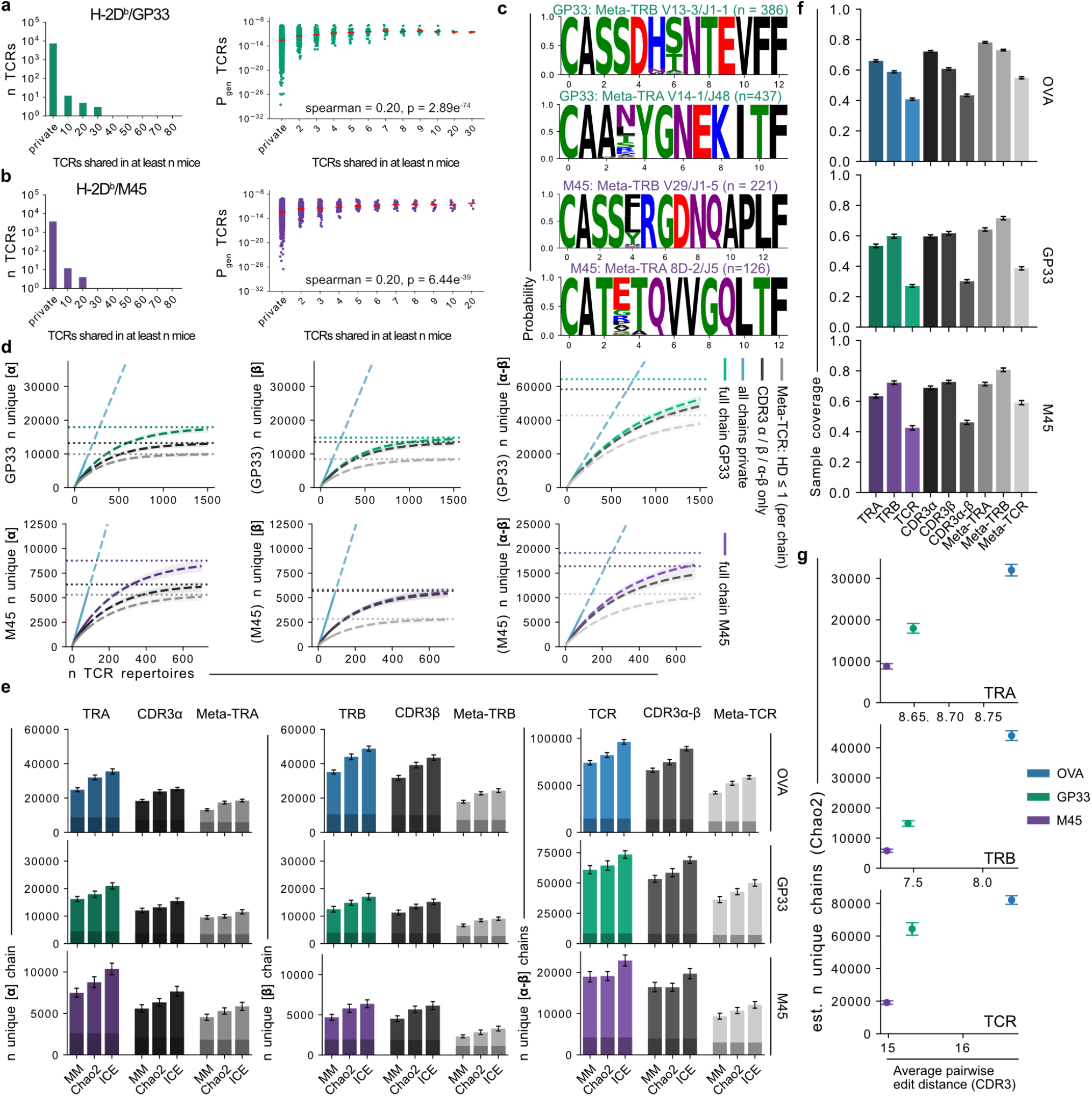
related to Figure 2: The total TCR solution space for functional epitope recognition is extremely large. (a) Left: Bar plot indicates TCR sharedness of GP33-reactive TCRs across donors (aa identity), right: Dot plot depicts correlation between TCR sharedness and TCR Pgen, line indicates median. (b), as in (A) but for M45 reactive TCRs. (c) CDR3*α* (lower) and CDR3*β* (upper) probability logo plots of Meta-TCRs within a hamming distance of 1. (d) Rarefaction and Extrapolation curves for estimated numbers of unique sequences for GP33-reactive (upper, green) and M45-reactive (lower, purple) sequences when sequencing a defined number of repertoires, colored tones depict full chains, dark gray CDR3, and light gray Meta-TCR. The cyan dashed line simulates the amount of expected individual sequences, if all sequences are private. From left to right: *α, β*, and paired *α*-*β*. HD denotes hamming distance. Solid line shows the rarefaction curve up to the observed numbers of repertoires and the dashed lines the extrapolated curve. Numbers calculated using iNEXT. Dotted line shows Chao2 asymptotic richness estimates. Shaded area depicts 95% confidence interval calculated using 100 bootstrap iterations. From left to right: *α, β*, and paired *α*-*β*. HD denotes hamming distance. (e) Estimated number of unique TCR chains or TCRs reactive against OVA (upper), GP33 (middle, green) or M45 (lower, purple) using Michaelis-Menten curve fitting (MM), Chao2, and Incidence Coverage-based Estimator (ICE) methods. Shaded areas depict observed numbers of sequences. (f) Estimated sample coverage for OVA, GP33, and M45 and all shown clone definitions. (g) Relationship between Chao2-estimated richness and average edit distance CDR3 within the set of epitope-reactive TCRs. All shaded areas and errorbars depict the 95% Confidence Interval calculated using 100 bootstrap samples calculated.

**Extended data 3.**
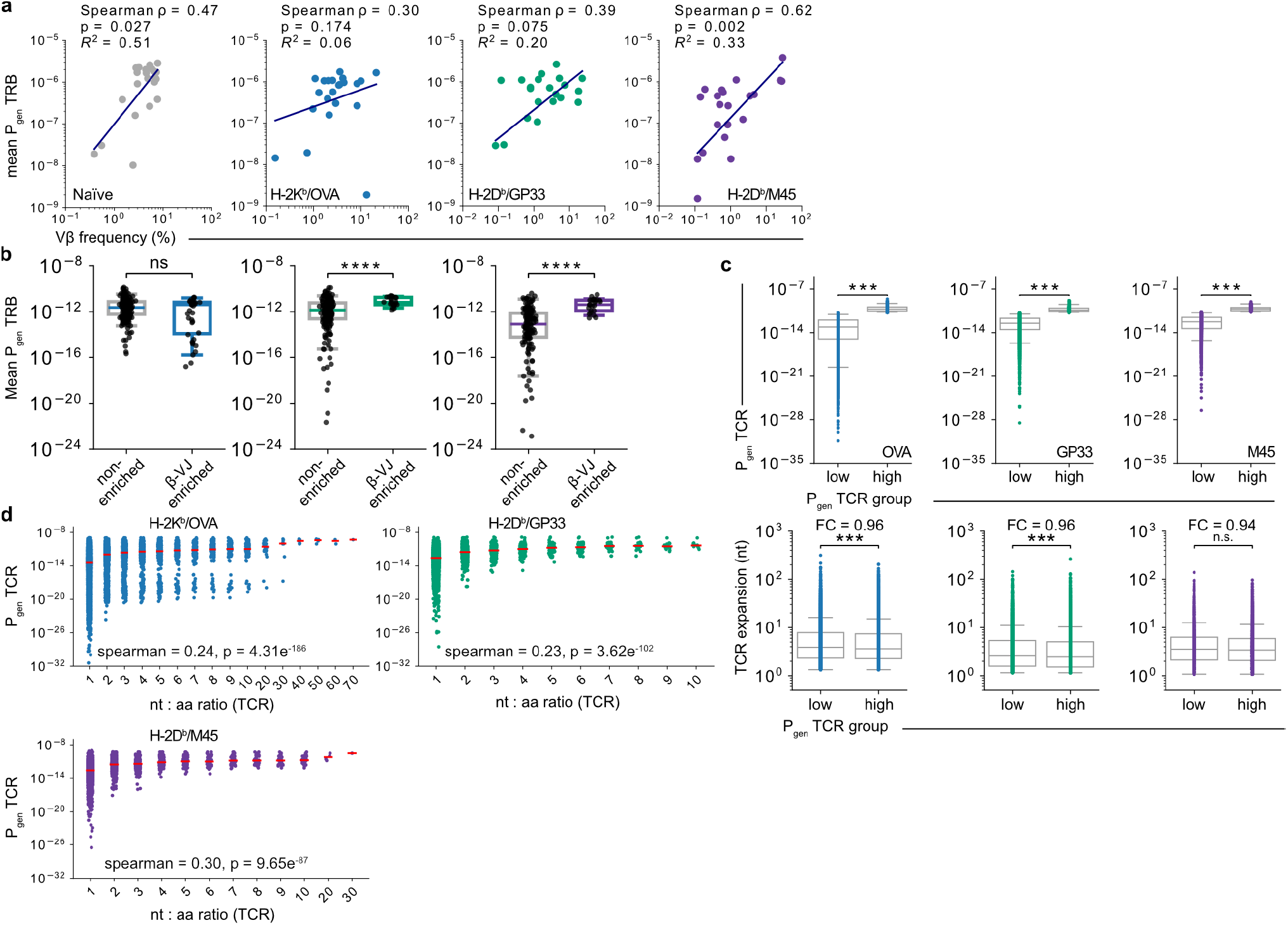
related to Figure 3: T cell recruitment is shaped by generation probabilities. (a) Scatterplots compare frequency of V*β* gene usage naïve, OVA, GP33 and M45 TCR libraries, against the average Pgen of the V*β* grouped TCRs. Each dot represents the average TCR Pgen within a unique V*β*. A linear regression was fitted on the data, and the Spearman Rank correlation was calculated. (b) Boxplots, compare average *β* P_gen_ per *β* V-J, each dot represents a unique *β* V-J. (c) Upper: Every TCR was split at the median distribution into a high or low P_*gen*_ group. Lower: Boxplots, compare TCR expansion (nucleotide identity) between TCR P_gen_ hi and low groups. (d) Scatterplots depict correlation between TCR P_gen_ and nt:aa ratio of clonotypes (aa identity) per epitope-reactivity. All statistical testing was done using a two-sided multiple Mann-Whitney test test with Holm-Šídák correction. * *p* < 0.05, ** *p* < 0.01, *** *p* < 0.001, **** *p* < 0.0001.

**Extended data 4.**
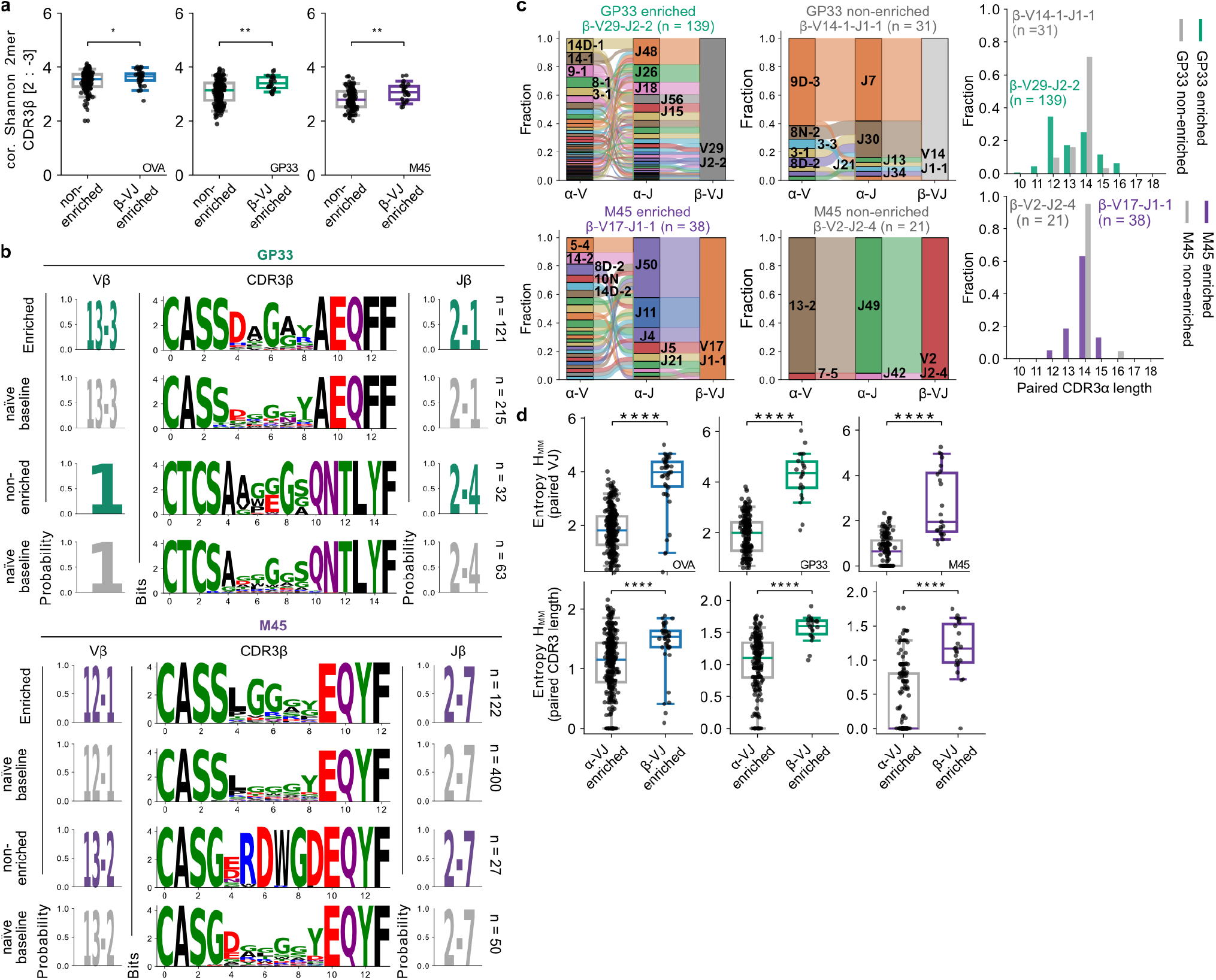
related to Figure 4: TCR germline architecture enables permissiveness in epitope-specific TCR features. (a) Left: *α* V-J *β* V-J ribbon plots of GP33- and M45-reactive TCRs. Displayed *β* V-J groups were selected for comparable size within IQR (0.35 - 0.65) of calculated HMM, right: CDR3*α* length spectratypes of the same *β* V-J group (gray, non-enriched, blue, enriched) (b) b) *α* V-J enrichment was calculated analogously as depicted for *β* V-J in Fig. 3d, for each *α* V-J group, HMM was calculated on *β* V-J pairing (as in Fig. 4a). Boxplots, compare HMM of the paired TCR germline (upper) and paired CDR3 length restriction (lower) (*α* V-J or *β* V-J, respectively) usage per enriched *α* V-J and enriched *β* V-J group. (c) Boxplots compare HMM of calculated kmers (*k* = 2, CDR3 C-term. trim = 2, N-term. trim = 3) per *β* V-J group between enriched and non-enriched, each dot represents a unique *β* V-J. (d) CDR3*β* logo plots of GP33- (left) and M45-(right) reactive TCRs in bits of enriched *β* V-J and non-enriched *β* V-J groups, alongside CDR3*β* logo plots of the same *β* V-J usage of the naïve baseline repertoire, left: probability logo of V*β*, right: probability logo of J*β*. All statistical testing was done using a two-sided multiple Mann-Whitney test with Holm-Šídák correction. * *p* < 0.05, ** *p* < 0.01, *** *p* < 0.001, **** *p* < 0.0001.

**Extended data 5.**
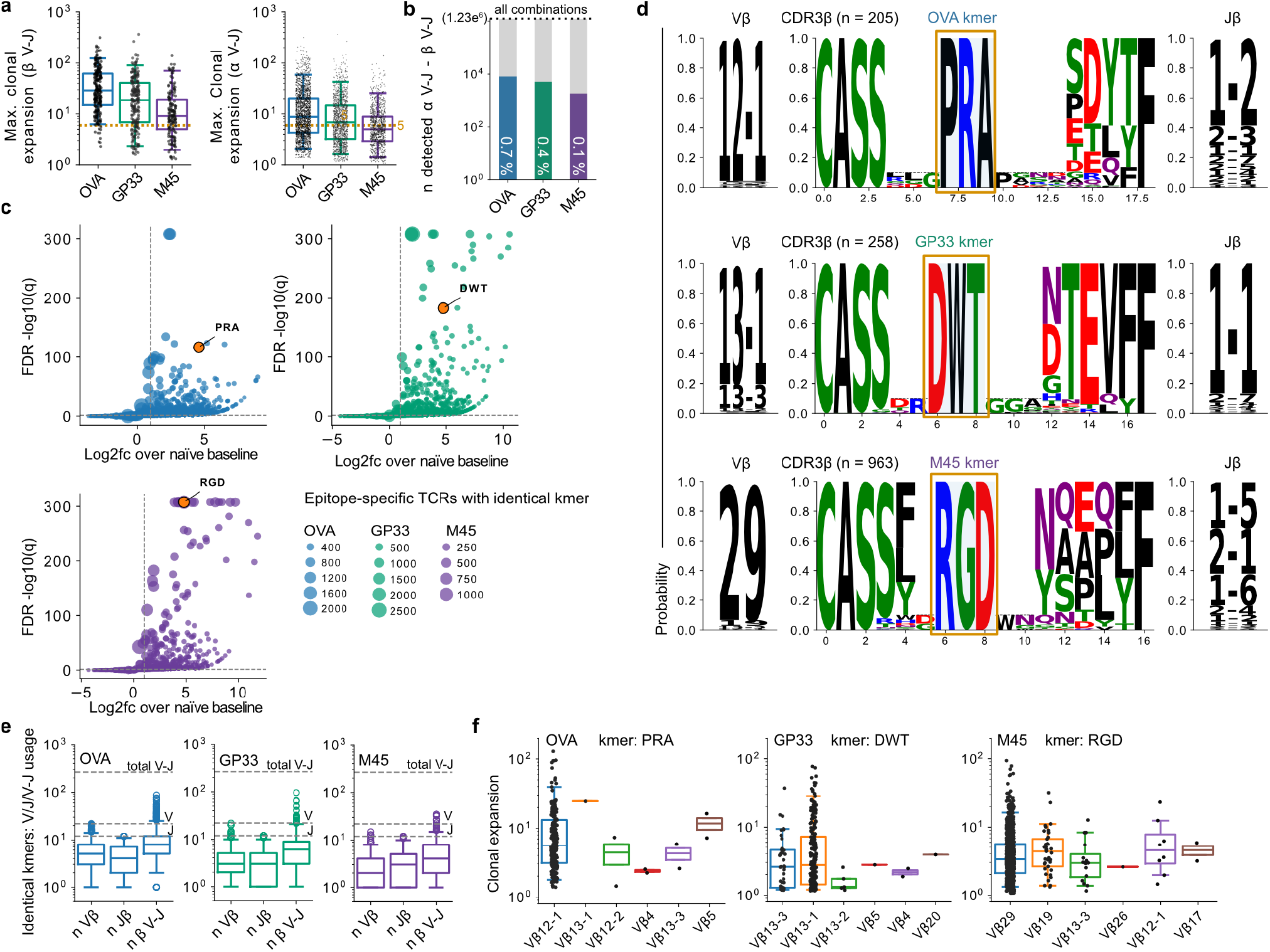
related to Figure 5: Almost all TCR germline architectures can give rise to a highly functional epitope-reactive TCR. (a) Maximal detected clonal expansion of a TCR per detected *β* V-J (left) and *α* V-J (right). Dotted line indicates expansion ≥5. (b) All detected TCR *α* V-J - *β* V-J gene segment combinations. Dotted line indicates all possible combinations. Numbers (white) indicate percentage detected of total combinations. (c) Kmer enrichment analysis across epitope-specificities, 3, 4, and 5mers were calculated over every CDR3*β* and compared to the naïve basline. Kmers were defined as enriched with a log2fc > 1 at q < 0.05. Each dot represents a unique kmer. (d) V*β*, CDR3*β* and J*β* probability logoplots of TCRs sharing significantly enriched kmers (orange box). All CDR3*β* sequences contribute to the logoplot, for multiple length alignment, gaps have been introduced around the fixed kmer, dashed line indicates that contributing amino acids at this position stem from TCRs below 10% of all contributing TCRs and the probability logo has been increased to 0.1 for better visbility. Plotted kmers have been selected for biggest contributing TCR size at a log2fc > 4. (e) Identified unique V*β*, J*β*, and *β* V-J of TCRs sharing identical CDR3*β* kmer motifs significantly enriched over naïve basline (log2fc > 1, q < 0.05), dashed lines indicate maximum of natural occuring unique V, J or V-J combinatorial segments. (f) Clonal expansion of TCR clonotypes sharing identical kmers shown in (d), seperated by their respective V*β* usage.

## 5 Methods

### 5.1 Mice

C57BL/6JOlaHsd (females, 6-10 weeks) were purchased from Inotiv. Mice were housed in a pathogen-free facility at the Technical University of Munich. All experimental protocols were approved by the standing committee for experimentation with laboratory animals of the Bavarian state authorities and performed in accordance with the corresponding guidelines (Upper Bavaria, Department 5 – Environment, Health and Consumer Protection; vote 55.2-2532.Vet_02-21-155).

### 5.2 Infection models and antigens

OVA_257-264_ (SIINFEKL) peptides were synthesized by peptides & elephants GmbH. Recombinant Listeria monocytogenes expressing SIINFEKL were kindly provided by Dietmar Zehn ^61^. Murine cytomegalovirus endogenously expressing M45_985-993_ (HGIRNASFI), and Lymphocytic choriomeningitis virus - Armstrong (LCMV) endogenously expressing GP33_33-41_ (KAVYNFATC), were kindly provided by Veit R. Buchholz. Mice were infected either intravenously (i.v.) with 2500 colony-forming units (CFU) of recombinant L.m.-SIINFEKL, or intraperitoneal (i.p.) with 2 × 10^5^ plaque forming units (PFU) LCMV or 500 PFU MCMV. The mice were sacrificed on day 8 post-infection (p.i.), and splenocytes were collected.

### 5.3 Preparation of single-cell suspensions

Spleens were passed through a 70 *μ*m cell strainer (Greiner BioOne), followed by gravity centrifugation (400*g*, 4 °C, 6 min). Spleen samples underwent erythrocyte lysis for 5 min at room temperature (RT) in 5 mL using 9 parts NH_4_Cl (0.17M, in ddH_2_O) with one part Tris-HCl (0.17 M, in ddH_2_O, pH = 7.5).

### 5.4 Multimer staining

Biotinylated pMHC molecules for the generation of pMHC multimers were refolded according to the protocol as previously described ^62^. 0.4 *μ*g of biotinylated pMHC class I molecule, 0.5 *μ*g of Streptavidin-APC and Streptavidin-BV421 and 50 *μ*l of FACS buffer for every 5 × 10^6^ cells, were preincubated for at least 30 min for multimerization. Cells were then incubated with the multimer mix for 40 min. 20 min before the end of the staining period antibodies for the staining of surface antigens were added. Propidium iodide for live/dead staining was added 5 min before the end of the staining period.

### 5.5 Antigen-specific activation for single-cell RNA and TCR sequencing

Splenocytes were harvested from L.m.-SIINFEKL-infected C57BL/6 mice and single-cell suspensions were prepared. 4 × 10^6^ splenocytes from each donor were separately incubated with 10^−4^ M SIINFEKL-peptide (dissolved in ddH_2_O) and 5 *μ*l of CD107a antibody in complete Roswell Park Memorial Institute 1640 medium (cRPMI), supplemented with 10% FCS, 0.025% L-Glutamine, 0.1% HEPES, 0.001% gentamicin and 0.002% streptomycin. CD107a (or LAMP-1) is a transmembrane protein found, among others, in cytotoxic granules of T cells (Terasawa2016). Degranulation of T cells leads to an accumulation of LAMP-1 molecules on the outer cell membrane and can thus be used as a marker for activated, cytotoxic T cells after antigen-specific stimulation (Betts2003). Samples were then further incubated at 37 °C, 5 % CO_2_ for 4 hours. Afterwards, the cells were washed with FACS buffer and stored on ice for further processing.

### 5.6 Fluorescent-activated cell sorting for processing on the 10X Genomics platform

Splenocytes from individual donors were stained for Fc-block (1:400) followed by TotalSeq™C anti-mouse Hashtag antibodies targeting MHC class I and CD45 according to the manufacturer’s instructions. Staining with fluorescent antibodies was performed in parallel. Splenocytes were stained for CD4, CD19, CD11c, NK1.1, CD8 and CD44. Individual donors were labelled with a unique TotalSeq™-C anti-mouse Hashtag antibody. 20 previously barcoded donors were pooled and in total 45,000 cells were sorted for CD4^−^ CD19^−^ CD11c^−^ NK1.1^−^ CD8^+^ CD44^hi^ and CD107a^+^.

### 5.7 pMHC multimer-dependent isolation

Splenocytes were harvested from infected (L.m.-SIINFEKL, MCMV, LCMV) C57BL/6 mice and single-cell suspensions were prepared. 4 × 10^6^ splenocytes from each mouse were stained for Fc-block (1:400) followed by pMHC-multimer staining (Streptavidin-APC and Streptavidin-BV421) as described. Afterwards, splenocytes were stained for TotalSeq™-C anti-mouse Hashtag antibodies targeting MHC class I and CD45 according to the manufacturer’s instructions. Staining with fluorescent antibodies was performed in parallel. Splenocytes were stained for CD4, CD19, CD11c, NK1.1, CD8 and CD44. Individual donors were labelled with a unique TotalSeq™-C anti-mouse Hashtag antibody. 20 previously barcoded donors were pooled and in total 45,000 cells were sorted for CD4^−^, CD19^−^, CD11c^−^, NK1.1^−^, CD8^+^, CD44^hi^, Streptavidin-APC^+^ and Streptavidin-BV421^+^.

### 5.8 Single-cell RNA and TCR sequencing (10X Genomics)

After cells have been sorted, they were centrifuged and the supernatant was carefully removed. Cells were resuspended in the Mastermix + 37.8 *μ*l of water before 70 *μ*l of the cell suspension were transferred to the chip. (Step 1.1 and 1.2 of the original protocol). After each step, the integrity of the pellet was checked under the microscope to ensure that all cells are loaded onto the chip. From here on, 10x experiments have been performed according to the manufacturer’s protocol (Chromium Next GEM Single Cell VDJ V1.1 with Feature Barcode, Rev D). QC has been performed with a high-sensitivity DNA Kit on a Bioanalyzer 2100 as recommended in the protocol and libraries were quantified with the Qubit dsDNA hs assay kit. All steps have been performed using RPT filter tips and DNA LoBind tubes.

### 5.9 Fluorescence-activated cell sorting for processing by Parse Bioscience platform

Splenocytes were harvested from infected (L.m.-SIINFEKL, MCMV, LCMV) C57BL/6 mice and single-cell suspensions were prepared. 4 × 10^6^ splenocytes from each mouse were stained for Fc-block (1:400) followed by pMHC-multimer staining (Streptavidin-APC and Streptavidin-BV421) as described. Afterwards, splenocytes were stained for CD4, CD19, CD11c, NK1.1, CD8 and CD44. 15.000 cells from each mouse were sorted for CD4^−^, CD19^−^, CD11c^−^, NK1.1^−^, CD8^+^, CD44^hi^, Streptavidin-APC^+^ and Streptavidin-BV421^+^ in separate wells, fixed and frozen according to the manufacturers instructions (Evercode Low Input cell fixation kit).

### 5.10 Single-cell RNA and TCR sequencing (Parse Bioscience)

After cells have been sorted, they were centrifuged at 400 x g, 4°C for 7 min and the supernatant was carefully removed. Cells were resuspended in 65 *μ*l of the Prefixation mastermix. 25 *μ*l of the fixation mastermix was added and incubated for 10 min on ice. Afterwards 8 *μ*l of the permeabilization solution was added and incubated for 3 min on ice. Then 110.4 *μ*l of the Stop-mastermix was added. The cells were placed in a Styrofoam box at room temperature and frozen at −80 °C. Before preparation of the sequencing libraries, the cells were thawed in a waterbath at 37°C and counted. From here on, single-cell RNAseq experiments were performed according to the manufacturer’s protocol (Evercode WT and TCR 100k). QC has been performed with a High sensitivity DNA Kit on a Bioanalyzer 2100 as recommended in the protocol and libraries were quantified with the Qubit dsDNA hs assay kit. All steps have been performed using RPT filter tips and DNA LoBind tubes.

### 5.11 Single-cell sequencing data processing

For 10X Genomics single-cell data, references GRCm39-2024-A, and GRCm38 were used for transcriptome and VDJ annotation (CellRanger 8.0.0), respectively. For Parse Biosciences singlecell data, annotation was performed via the Trailmaker cloud service (Parse Biosciences - Pipeline v1.6.1). Data preprocessing has been performed according to the current best practice in scRNA sequencing analysis. Data analysis was performed with Scanpy V1.8.2 and Scirpy 0.10.1. Briefly, cells with fewer than 200 genes as well as genes present in fewer than three cells were excluded. Counts were normalized per cell and log-transformed. The top 5000 highly variable genes were identified and filtered using Pearson residuals (Scanpy). The data were batch-corrected using batch-balanced k-nearest neighbors (bbknn) for individual experiments. DNA-Barcoded-demultiplexing and cell doublet detection were performed with Scrublet and HashSolo included in Scanpy, respectively.

### 5.12 TCR clonotype definition, reactivity annotation and size normalization

Clonotype analysis was performed using Scirpy ^63^. Cells belonging to one clonotype were defined to have identical TCR*α* and *β*-chain amino acid sequences (CDR3 amino acid sequence, as well as identical VJ-gene annotation), unless indicated as nucleotide identity (then: CDR3 nucleotide sequence, as well as identical VJ-gene annotation). Single cells that harbored more than a single TCR*α* or *β*-chain were excluded from further analysis. Antigen-reactive TCR clonotypes were defined experimentally through binding to a given pMHC multimer, or by expressing CD107a after antigenic stimulation, while being identified with a minimal clonal expansion of two in a single donor, after single-cell RNA and TCR sequencing. To compare clonal expansion between donors, size factor normalization was performed. The average single-cell count was calculated across all individuals; every single-cell count of every donor was divided by the average count, giving a size factor for every donor, respectively. The clonal expansion of each TCR was calculated within each donor and multiplied by the donor-specific size factor.

### 5.13 TCR cloning

DNA templates were designed in silico and synthesized by Twist Bioscience. DNA constructs for retroviral transduction had the following structure: Murine Kozac sequence followed by TCR*β*, followed by P2A, followed by TCR*α*, cloned into the pMP72 vector (kindly provided by Wolfgang Uckert, Berlin, Addgene plasmid backbone #108214).

### 5.14 Culture of cell lines

RD114 cell lines were grown in complete Dulbecco’s Modified Eagle Medium (cDMEM), supplemented with 10% FCS, 0.025% L-Glutamine, 0.1% HEPES, 0.001% gentamicin and 0.002% streptomycin. TCR^−^, human CD8^+^ Jurkat triple parameter reporter (JTPR) cell lines and primary T cells were cultured in cRPMI. Primary T-cell culture was additionally supplemented with 25 IU mL^−1^ IL-2. All cells were grown in a 37°C humidified, 5% CO_2_ incubator. JTPRs were originally obtained from Peter Steinberger (Medizinische Universität Wien).

### 5.15 Flow cytometry

Single-cell suspensions of spleens or JTPR were stained with the respective antibody panel for 25 min at 4 °C in the dark. After washing with FACS buffer (PBS with 0.5% BSA and 2 mM EDTA), the cells were stained with PI for 5 min and washed again. Data were collected by flow cytometry on a CytoFLEX S or CytoFLEX LX flow cytometer (Beckman Coulter). For analysis, FlowJo software (v10.10.0, FlowJo LLC) was used.

### 5.16 Virus production and transduction

For retrovirus production, RD114 packaging cells were transfected with the retroviral vectors (mp71, a kind gift from W. Uckert, added as Addgene plasmid backbone #108214) encoding for the respective SIINFEKL-reactive TCRs via calcium phosphate precipitation. The supernatant of the RD114 cells was collected at 72 h after transfection and purified from remaining cells by centrifugation at 600 x g. at 4 °C for 7 min. The supernatant was stored at 4 °C and used within 4 weeks after collection. Retroviral transduction of JTPR cells with the respective TCR was achieved by spinoculation. In brief, 400 *μ*l of the TCR-RD114 supernatant was centrifuged at 3,000*g* at 32 °C for 2 h in a tissue-culture-untreated 48-well plate coated with RetroNectin according to the manufacturer’s instructions. Afterwards, the supernatant was discarded, and JTPR cells were added at a final concentration of 50,000 cells per 400 *μ*l per well. Cells were then spinoculated at 800*g* at 32 °C for 20 minutes. After 3 days in culture, transduction efficacy was assessed by assessing co-expression of murine TCR*β* with human CD3 (hCD3) via flow cytometry. JTPR cells were purified for TCR expression by flourescence-activated cell sorting for mTCR*β*^+^ and hCD3^+^.

### 5.17 Antigen-specific activation of JTPR

TCR-transduced JTPR were purified for hCD3 and mTCR*β* expression (as a marker for a transgenic TCR) via flourescence-activated cell sorting before they were used in a stimulation assay. Spleens of naïve, wild type C57Bl/6 mice were harvested as antigen presenting cells and 1.2 × 10^5^ splenocytes were loaded with 10^−5^M of the SIINFEKL-peptide, a negative (cRPMI only) and a positive control (PMA 0.025 *μ*g ml^−1^, Ionomycin 1.0 *μ*g ml^−1^). 2.4 × 10^4^ TCR transduced JTPR were added to each condition in parallel in separate stimulations. Samples were then incubated at 37 °C for 24 hours. The percentage of NFAT^+^ (GFP reporter) JTPR after stimulation was assessed via flow cytometry. Live/dead discrimination was performed with propidium iodide.

### 5.18 Calculation of Miller-Madow normalized Shannon entropy and Gini coefficient

For all TCR*α* and/or *β* analyses, Shannon entropy was estimated from the observed categorical frequencies using natural logarithms. For a distribution containing K observed categories and n total observations, the empirical entropy was calculated as

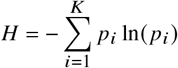

where *p*_*i*_ = *n*_*i*_ */ n* is the observed relative frequency of category i. Categories with zero frequency were not included. To avoid underestimation of entropy in small samples, the Miller-Madow correction was applied.

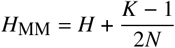

where K is the number of observed categories and *N* is the number of non-missing observations contributing to the distribution. The correction is zero when only one category is observed.

Separately, for calculation of the CDR3*β* position flexibility ratio, epitope-reactive and naive background CDR3 sequences were compared within matching length and *β* V-J combination. Amino-acid frequencies were calculated independently at every CDR3 position. For every position j, the epitope-specific corrected entropy was calculated as H_MM_ and normalized to the maximum amino acid variability (ln20).

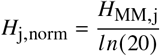

where zero represents complete conservation of a single amino acid. The CDR3*β* position flexibility ratio was defined as the mean normalized positional entropy of the epitope-specific repertoire divided by the corresponding mean background entropy. For each V-J combination, we then computed the mean positional entropy across all CDR3 lengths, weighting each length group by the number of contributing TCR sequences. This weighted mean entropy was normalized to that of the corresponding V-J combination in the naïve repertoire, yielding a CDR3*β* positional flexibility ratio. Ratios below one indicate reduced positional variability in amino acid usage relative to the naïve baseline.

In addition we quantified inequality within a distribution of V*α* and V*β* gene segments. The Gini coefficient was calculated as

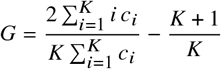

Where the counts *c* of every category *i* are determined and ordered in ascending order, and *K* determines the count of unique observed categories. The Gini coefficient was calculated for V*α* and V*β* in both the epitope-reactive repertoires and naïve repertoires per mouse. Gini coefficient of naïve mice were averaged and subtracted from the Gini coefficient of every epitope-reactive repertoire for V*α* and V*β*, respectively.

### 5.19 Epitope-specific prediction model

For each epitope, the pretrained ESM-2 protein language model was fitted with a fully connected classification head and finetuned with LoRA ^64^ to discriminate between reactive and unreactive TCRs. The model received the full, concatenated *α*- and *β*-TCR-chains, with the constant region truncated, from the initial 10x experiments as input. Sequences with a CDR3 lengths greater than 30 amino acids or detected for two specificities were excluded, resulting in n_OVA_=6,880, n_GP33_=1,824, n_M45_=1,118 positive sequences which were complemented by n_naïve_=50,404 negative TCRs from five uninfected C57BL/6 mice at a ratio of 1:5.13. After adding start, end, and separating tokens, the end-padded sequences were tokenized as model inputs. To prevent data leakage, this dataset was split into six folds based on identical CDR3*α* or CDR3*β* clonotype groups, separated through Leiden clustering ^65^ with one fold exclusively held out for final testing. Models were trained in a five-fold cross-validation on the remaining folds via the binary cross-entropy loss on maximal batch size fitting in memory until early stopping was reached on the validation AUC. For each fold, hyperparameters were optimized with Optuna ^66^ for 48 hours on a single NVIDIA A100 GPU (80GB). For TCR classification, a threshold of 0.5 was applied.

### 5.20 Estimation of the total number of unique, epitope-reactive TCRs

Chao2 is an incidence-based estimator that is particularly well suited to our dataset, as it infers diversity from the presence or absence of individual sequences across a large number of independently sampled immune repertoires. Chao2 and its abundance-based counterpart, Chao1, have previously been used to estimate TCR richness across a range of biological settings, including human peripheral blood, the thymus, and infectious disease ^33–35^. More recently these estimators have also been benchmarked for TCR repertoire diversity profiling ^36^. The iNEXT R package ^67^ was used to calculate the sample-size-based rarefaction and extrapolation curves, as well as the Chao2 estimate with 95% confidence interval using 100 bootstrap repetitions, and the sample coverage. Three different methods were employed to estimate the total number of unique, epitope-reactive TCRs: Chao2, ICE, and Michaelis-Menten curve fitting. The bias-corrected Chao2 estimate is calculated by ^68^:

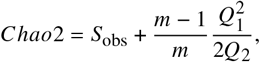

where *S*_*obs*_is the observed number of unique sequences found in *m* repertoires, *Q*_*1*_ the number of sequences found in a single repertoire, and *Q*_*2*_ is the number of sequences found in two repertoires. The Incidence coverage-based (ICE) estimate is calculated by ^37^:

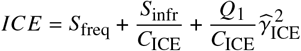

where *S*_*freq*_ and *S*_*infr*_ are the counts of frequent and infrequent unique sequences, respectively. Using the definition in the original paper, sequences found in more than 10 mouse repertoires were considered frequent, while those found in less than or equal to 10 were defined as infrequent. *Q*_*1*_ is the number of sequences found in exactly one repertoire. *C*_*ICE*_ is the sample coverage estimate and 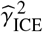 is the estimated coefficient of variation.

Following Schober _38_, we fitted the rarefaction to the Michaelis-Menten curve of shape

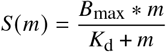

by estimating the parameters half-life *K*_*d*_ and limiting rate *B*_*max*_ to minimize the residual sum of squares. *m* depicts the number of analyzed mouse repertoires. The limiting rate *B*_*max*_ is used as the asymptotic estimate of the saturation.

### 5.21 Construction of meta-TCRs

To aggregate similar TCRs within the same V-J germline and CDR3 profile, we calculated meta TCRs. First, the pairwise Hamming distance of all CDR3-*α* and *β* sequences were calculated. Iteratively, the sequence with the highest frequency was chosen as a seed and absorbed its neighbors within Hamming distance 1 and removed from the set. This process was repeated until all sequences were assigned to a meta TCR seed.

### 5.22 Kmer CDR3 motif enrichment

Local CDR3 motif enrichment was assessed using an approach adapted from the local-convergence framework introduced in GLIPH and GLIPH2^17,69^. Local CDR3 motif enrichment was assessed by comparing epitope-specific TCR sequences with the naïve repertoire baseline. Terminal residues of full-length CDR3 amino-acid sequences were trimmed (two N-terminal and three C-terminal residues) to reduce contributions from germline-encoded V- and J-derived sequence. All contiguous 3–5 amino-acid motifs were enumerated, and motif prevalence was calculated as the fraction of unique CDR3 sequences containing each motif. Motifs represented by at least three epitope-specific CDR3s were tested for enrichment relative to the complete naïve repertoire using a one-sided Fisher’s exact test, followed by Benjamini–Hochberg correction for multiple testing. Fold enrichment was calculated from motif frequencies in the epitope-specific and naïve repertoires. To reduce redundancy among nested k-mers, motifs occurring in identical sets of epitope-specific CDR3 sequences were collapsed, retaining the shortest representative motif.

